# Direct interaction between cancer cells and fibroblasts promotes early chemoresistance to standard-of-care drug therapy in small cell lung cancer (SCLC)

**DOI:** 10.64898/2026.04.27.721078

**Authors:** Nilu Dhungel, Brian Latimer, David Custis, Ana-Maria Dragoi

## Abstract

Cancer-associated fibroblasts (CAFs) are now recognized as key regulators of tumor progression and therapeutic resistance, yet the cancer cells-fibroblasts crosstalk that ultimately promotes chemoresistance remain incompletely understood. Here, we show that direct physical contact between small cell lung cancer (SCLC) cells and lung fibroblasts induces early resistance to standard-of-care chemotherapeutic agents, etoposide and cisplatin. Using both 2D and 3D co-culture models, we demonstrate that early acquired therapy resistance entails cell-cell contact and cannot be recapitulated by conditioned media alone. Mechanistically, direct interaction promotes transcriptional reprogramming in cancer cells, including upregulation of YAP1 and epithelial-to-mesenchymal transcription factors (EMT-TFs), which partially mediate the resistant phenotype. A high-throughput drug screening identified idarubicin as a compound that retains efficacy despite fibroblast-mediated protection, suggesting it could bypass microenvironment-induced resistance early on. Together, our findings identify direct tumor–fibroblasts contact as an early driver of chemoresistance and highlight a potential therapeutic strategy targeting cell-cell interactions within the tumor microenvironment.

## Introduction

Despite remarkable advances in diagnosis and treatment, cancer remains one of the leading causes of mortality worldwide, with nearly 10 million deaths in 2022 [1, 2]. Among the various cancer treatments available for cancer management-including surgery, gene therapy, immunotherapy, radiation therapy, and laser therapy - chemotherapy is the most common and promising treatment [3]. While significant progress has been made since the introduction of chemotherapy in the 1900s, chemoresistance, the capacity of cancer cells to resist or adapt to a therapeutic agent, continues to be a significant challenge in cancer therapy, limiting treatment efficacy, driving relapse, and contributing to poor patient survival [3, 4]. Studies show that nearly all chemotherapeutic drugs could ultimately encounter resistance, even after an initial favorable response [5]. Chemoresistance may manifest as either intrinsic, inherent to the cancer cells, or acquired, arising over the course of treatment [3]. Regardless of its origin, chemoresistance leads to disease relapse and metastasis, significantly hindering efforts to improve the clinical outcomes of cancer patients [4, 6].

The molecular mechanisms of chemoresistance occur at multiple levels and include drug efflux pumps, mitochondrial alterations, enhanced DNA repair, epithelial-to-mesenchymal transition (EMT), activation of survival pathways, and inactivation of tumor suppressor genes [1, 3, 6, 7]. Overexpression of ATP-binding cassette (ABC) transporter proteins in various cancer types, including leukemia, neuroblastomas, ovarian, and breast cancers enables them to develop resistance to therapeutic agents [6, 8]. In addition to alterations in the efflux pumps, defects in DNA repair mechanisms enable cancer cells to be more resistant to chemotherapeutic agents. Studies have shown that tumor cells with increased nucleotide excision repair (NER), a major pathway for regulating DNA repair, have a higher capacity to repair DNA damage induced by chemotherapeutic drugs, leading to treatment resistance [9]. For example, overexpression of NER is linked to resistance to platinum-based chemotherapy treatments in ovarian cancer [9, 10]. Furthermore, inhibition of the NER pathway significantly increased the sensitivity of tumor cells to cisplatin [9, 11]. Inhibition of pro-apoptotic pathways or upregulation of anti-apoptotic pathways also induces resistance to chemotherapy in cancer cells [12, 13]. Crucial tumor suppressors, such as p53, are often mutated or inactivated in cancer cells, thereby compromising their ability to promote apoptosis [14]. In addition, multiple studies have also shown that various signaling pathways involved in maintaining stemness in cancer cells also contribute to the induction of chemoresistance [15].

Recently, the tumor microenvironment (TME), composed of various cellular and non-cellular components, has been shown to play a critical role in drug resistance. Increasing evidence supports the notion that the microenvironment is a key driver of cancer relapse and treatment failure across various cancers [16, 17]. Studies have shown that the tumor microenvironment diminishes the efficacy of cancer treatments and enhances chemotherapy resistance by fostering aberrant vasculature, hypoxia, a rigid matrix, and altered paracrine factor composition [4]. Stromal proportions of the tumor are determinants of the therapeutic success in various cancer types, including glioblastoma, prostate, colorectal, breast, pancreatic, cervical, and ovarian cancers [17]. It has been shown that tumors resistant to alkylating agents *in vivo* become sensitive to the same agents when cultured outside the body in the absence of their TME, indicating that stromal cells may contribute to the development of resistance to anticancer drugs [18, 19].

Cancer-associated fibroblasts (CAFs), a major cellular component of the TME, mediate a *de novo* chemoresistance, facilitating cancer cells’ protection from apoptosis induced by chemotherapy, radiotherapy, or receptor-mediated cell death [17, 20]. CAFs can induce either soluble factor-mediated drug resistance (SFM-DR) or cell adhesion-mediated drug resistance (CAM-DR) [20]. SFM-DR is induced by CAF-secreted cytokines, chemokines, growth factors, and extracellular vesicles, which activate pro-survival pathways in cancer cells, thereby reducing drug sensitivity. CAM-DR, on the other hand, is mediated by tumor cell integrins binding to stromal fibroblasts or to components of the extracellular matrix (ECM), such as laminin, collagen, and fibronectin, which participate in cell adhesion [17, 20]. Studies have demonstrated that CAFs can promote chemoresistance through multiple mechanisms, including the regulation of drug efflux pumps, mitochondrial reprogramming, enhanced DNA repair, induction of epithelial-to-mesenchymal transition (EMT), activation of pro-survival signaling pathways, and inhibition of tumor suppressor function via the release of exosomes and various secreted factors [17]. While the contribution of CAF-secreted factors to chemoresistance is well established [21-25], the role of direct physical interactions between cancer cells and fibroblasts is less understood.

At the same time, the simultaneous occurrence of EMT and chemoresistance has drawn attention to the idea that EMT induction may contribute to drug resistance. Previous studies show that drug-resistant tumor cells exhibit EMT features and that inhibiting EMT restores their drug sensitivity [26]. A previous study demonstrated that an adriamycin-resistant MCF-7 cell line and a vinblastine-resistant ZR-75-B cell line exhibit the phenotypic characteristics of EMT, including a significant increase in vimentin expression and a reduction in desmosome and tight junction formation [27, 28]. There is also considerable overlap between the signaling pathways involved in EMT and chemoresistance, with EMT-inducing signals directly linked to treatment resistance in cancer [29]. Activation of various cellular signaling pathways, such as Wnt, Notch, TGF, and Hedgehog, which are critical for inducing EMT, has been shown to contribute to chemoresistance across multiple cancer types [29]. TGF-β, a key signaling pathway in EMT induction, plays a critical role in chemoresistance, and studies have shown that treatment with TGF-β antibodies increased the drug sensitivity of the alkylating agent-resistant tumors to drugs such as cisplatin and cyclophosphamide [28, 30]. Likewise, overexpression of Wnt3, another critical signaling pathway in EMT induction, in human epidermal growth factor receptor 2 (HER2)-overexpressing breast cancer cells promotes a partial EMT-like transition by activating the Wnt/β-Catenin pathway, resulting in trastuzumab resistance. Moreover, siRNA-mediated knockdown of Wnt3 reduces EMT markers and decreases cancer cell invasiveness [31]. In addition to signaling pathways, EMT transcription factors (EMT-TFs), the master regulators of EMT, can directly contribute to treatment resistance in malignancies [29]. Various EMT-TFs, including Snail, Twist, and FOXC2, drive the overexpression of ABC transporters, which contribute to treatment resistance by facilitating the efflux of cytotoxic drugs from malignant cells, lowering intracellular drug concentrations, and reducing therapeutic efficacy [29, 32]. For example, ZEB1, Snail and Slug, key EMT-TFs, are associated with poor response to temozolomide in glioblastoma patients [33] and cisplatin resistance in ovarian cancer, respectively [34]. Furthermore, cancer cells undergoing EMT often exhibit stem-like properties, as there are remarkable similarities in pathways that drive EMT and those that regulate CSCs [28]. These pathways arm cancer cells with self-renewal capacity, enhanced survival, and increased resistance to therapy. Thus, the co-expression of key stemness-associated transcription factors, OCT4 and Nanog, promotes CSC properties and confers gefitinib resistance in NSCLC by regulating the Wnt/β-catenin signaling pathway [35].

Small cell lung carcinoma (SCLC) is a highly aggressive form of lung cancer that is characterized by early dissemination, extremely poor prognosis, a good initial response to chemotherapy, and high frequency of metastasis at autopsy [36-39]. Although SCLC responds well to platinum-based chemotherapy with an initial response rate of greater than 60% to 70%, nearly all patients ultimately develop rapid acquired resistance upon relapse [40, 41]. In addition to tumor intrinsic mechanisms of resistance, including altered DNA damage repair, altered differentiation state, upregulation of drug resistance proteins, and altered tumor metabolism, tumor extrinsic factors, such as alterations in the tumor microenvironment, play a critical role in inducing therapy resistance in SCLC. Fridman *et al*. showed that SCLC develops increased therapy resistance in the presence of basement membrane components such as laminin [40, 42]. Inhibition of cellular interaction with the extracellular matrix by targeting β1 integrins or using a tyrosine kinase inhibitor restores SCLC responsiveness to etoposide to levels observed in treatment-naïve cells [40] suggesting the roles the ECM plays in inducing chemoresistance in SCLC. A recent study by Lu *et al*. examined the role of cancer-associated fibroblasts in dynamic phenotypic reprogramming in SCLC. The authors demonstrated that transwell indirect co-culture of SCLC with fibroblasts not only drove phenotypic reprogramming of SCLC but also promoted a highly inflamed tumor microenvironment. Additionally, they showed that the SCLC in indirect contact with fibroblasts were more resistant to cisplatin and etoposide compared to the monocultured cells, suggesting that SCLC tumors with high CAFs infiltration could exhibit increased chemotherapy resistance [43]. Inhibition of CAF-activated JAK/STAT3 signaling by using JAK1/2/3 inhibitors increased the sensitivity of the drugs [43] further reinforcing the critical role of CAFs in mediating therapy resistance in SCLC.

We have previously demonstrated, using an *in vitro* SCLC and lung fibroblast model, that direct interaction between cancer cells and fibroblasts leads to profound early reprogramming of cancer cells, resulting in the development of a hybrid, reversible EMT phenotype [44, 45]. We observed the upregulation of a select group of EMT-TFs, ZEB1, ZEB2, and TWIST2. Additionally, there was significant upregulation of Yes-associated protein 1 (YAP1), a core component of the Hippo/YAP pathway, in cancer cells in direct contact with fibroblasts, but not in indirect contact. Our previous results revealed the significance of direct contact between cancer cells and fibroblasts in inducing a hybrid and transient EMT phenotype in cancer cells. Using the same model, we extended our investigation to explore the role the direct physical interaction between cancer cells and fibroblasts plays in chemoresistance. We discovered that direct interaction-induced EMT leads to early chemoresistance, partially via the YAP1 pathway. We further demonstrate that inhibition of YAP1 with verteporfin resensitizes cells to therapeutic treatments, even when they are in contact with fibroblasts. Using a 3D model of interaction, we performed a high-throughput drug screening using live-cell imaging and the NIH Clinical Collections drugs to uncover new potential treatments that are efficient against SCLC when in direct contact with fibroblasts. We identified idarubicin as a chemotherapeutic agent capable of targeting both SCLC and CAFs when the cells are in direct contact.

## Results

### Drug dose optimization in SCLC and lung fibroblast cell lines

To understand better the role of interaction between cancer cells and fibroblasts in chemoresistance, we used a model of SCLC lines (H69 or H209) and lung fibroblasts (CCD8s) cultured in direct or indirect contact [44]. The standard-of-care drugs for SCLC, cisplatin and etoposide [46-48], were used as chemotherapeutic agents in our studies. First, to identify the doses of drugs that induce apoptosis in cancer cells but not in fibroblasts, we performed a drug-optimization experiment. We treated cancer cells (H209 and H69) and fibroblasts (CCD8) with various serial dilutions of cisplatin alone or etoposide alone. The three cell lines were treated separately with serial dilution concentrations (50 μM, 25 μM, 12.5 μM, 6.25 μM, and 3.125 μM) of the two chemotherapy agents. We used the IncuCyte® Annexin V NIR Dye to label apoptotic cells and acquired live microscopy using the IncuCyte S3 microscope (Figure 1A). Annexin V red-positive cells were identified as apoptotic, while red-negative cells were identified as live cells using the integrated software of IncuCyte S3 microscope. We observed that both cisplatin and etoposide induce significant cancer cell death at 12.5 and 6.25 μM after 72h, without causing cell death in fibroblasts (Figure 1B and 1C, left panels and Supplemental Fig 1). We also quantified differences between conditions by calculating the area under the curve (AUC) and used these values to assess statistical significance (Figure 1B and 1C, right panels). We further verified these results by Western blot using cleaved and uncleaved caspase 3 and caspase 7 as readouts in cells treated with etoposide for 72h (Figure 1D). Consistent with our live-imaging data, Western blot analysis showed that SCLC cells are sensitive to lower doses of etoposide, whereas CCD8 fibroblasts are not. The optimization experiments also revealed that the H209 SCLC cells are more sensitive to etoposide compared to cisplatin. We also noticed that while both SCLC cell lines follow a similar pattern, H209 cells are slightly more sensitive to the same doses of chemotherapeutic agents than H69 cells. We, therefore, used H209 for further chemotherapy response characterization.

**Figure 1.**
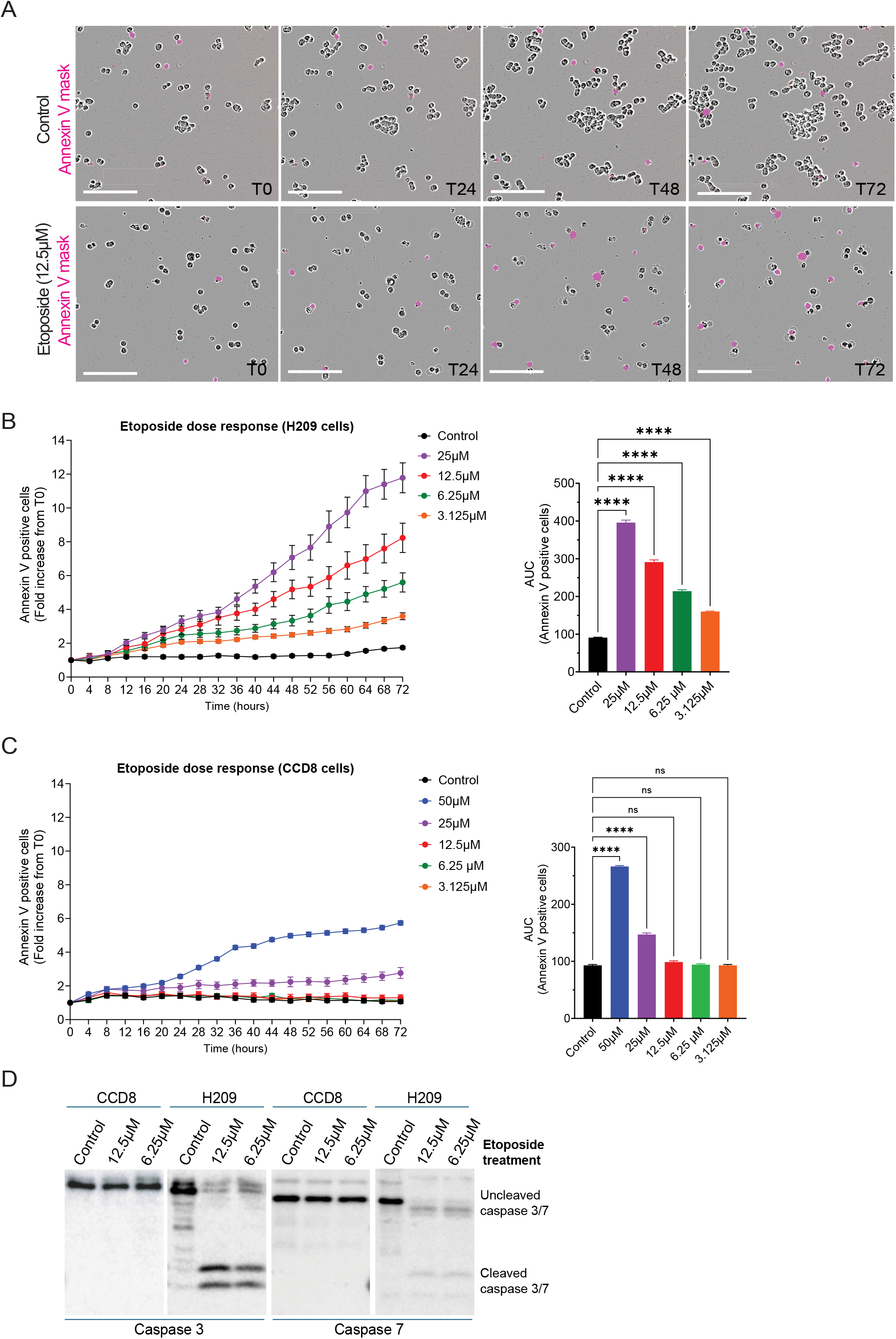
Drug optimization and apoptosis assessment in SCLC cancer cells. **(A)** Representative images of Control and Etoposide-treated H209 cells acquired using the IncuCyte S3 live cell imaging system. Annexin V red dye was used to identify apoptotic cells. The pink mask represents the annexin V-positive apoptotic cells. Bar = 200 µm. **(B-C)** Etoposide dose response analysis in **(B)** H209 cancer cells and **(C)** CCD8 fibroblasts. Left panels show the time course increase in apoptotic cells number identified as annexin V-positive cells using IncuCyte S3 live microscopy. Graphs are presented as fold increase from T0 using the “Red Area / Phase Area Normalized to T0” software analysis. Right panels show “Area under the curve” (AUC) analysis using the Prism software. AUC was computed from “Fold increase annexin V-positive cells” *vs* time using the trapezoidal rule across 72 hours. Statistical significance between Control and treatment groups was assessed using one-way ANOVA with Dunnett’s post hoc correction. Data are shown as mean ± SEM from one representative experiment (6 replicates/plate). p-values are indicated in the graphs: ns, not significant; *\*\*\*\**, p< 0.0001. **(D)** Immunoblot analysis of cleaved and uncleaved caspase-3 and caspase-7 in H209 cancer cells and CCD8 fibroblasts after etoposide treatment for 3 days.

### Direct contact between cancer cells and fibroblasts induces chemoresistance in cancer cells

Having established the optimal drug doses that induce apoptosis in cancer cells but not in fibroblasts, we next tested whether cancer cells in direct contact with fibroblasts develop resistance to chemotherapy during interaction. For this, we labeled only the H209 cancer cells with a green fluorescent dye and cultured them alone or in direct contact with unlabeled fibroblasts for 24h prior to treatment. After the initial culture, cells in both conditions were treated with etoposide or cisplatin at 6.25 μM or 12.5 μM for an additional 72h. We used IncuCyte® Annexin V NIR Dye to label apoptotic cells and acquired live-cell imaging using the IncuCyte S3 microscope. Double-positive cells (red and green) were identified as apoptotic, while single-positive cells (green only) were identified as live cancer cells using the integrated software of IncuCyte S3 microscope (Figure 2A). Cell death was quantified as the ratio of red and green cells to total green cells, and as the fold increase over T0 at the start of acquisition. The dose-response time-course graph showed that the number of apoptotic cancer cells is lower when cancer cells are grown in direct contact with fibroblasts than when they are grown alone at the same dose of etoposide (Figure 2B, left panel). As before, we quantified differences between conditions by AUC and used these values to assess statistical significance (Figure 2B, right panel). We also performed Western blot analysis of cancer cells after treatment and FACS sorting to assess the levels of cleaved and uncleaved caspase 7, in order to validate the microscopy data. Consistent with the live microscopy data, the Western blot analysis of H209 cells treated alone or in direct contact with CCD8 cells showed that cancer cells acquire therapy resistance over 96h in contact with fibroblasts (Figure 2C). Altogether, these results suggest that cancer cells in direct contact with fibroblasts in the microenvironment become less sensitive to chemotherapy treatment.

**Figure 2.**
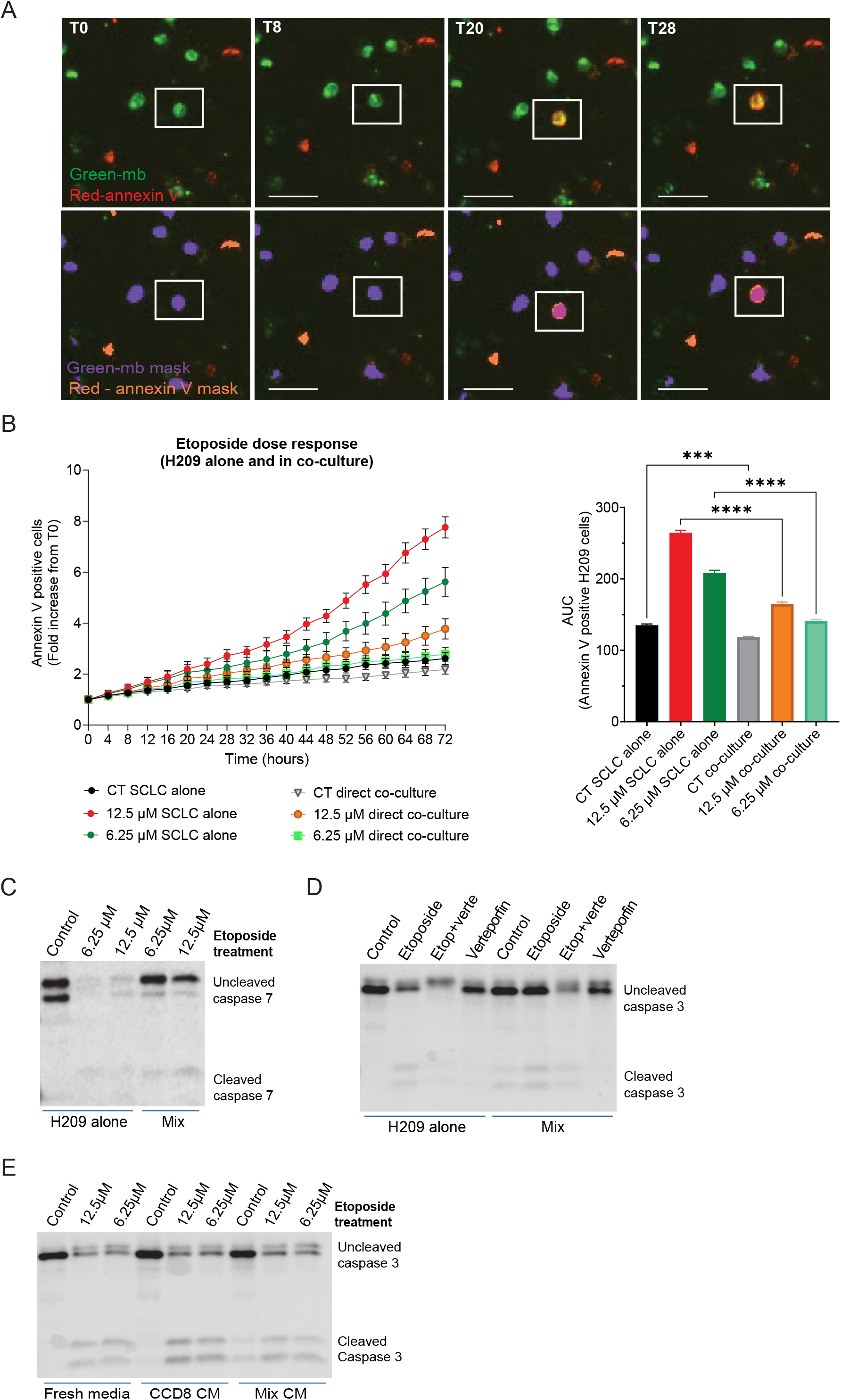
Direct interaction between SCLC cancer cells and lung fibroblasts induces chemoresistance to etoposide treatment. **(A)** Representative images acquired using the IncuCyte S3 live cell imaging system at different time points after treatment with etoposide in cancer cells-fibroblasts co-culture. Cancer cells only were labeled with a green fluorescent dye prior to co-culture. Annexin V red dye was used to identify apoptotic cells. The purple mask identifies live green cancer cells, while the orange mask identifies Annexin V-positive apoptotic cells. Cells positive for both green and red signals represent apoptotic cancer cells. Bar = 50 µm. **(B)** Etoposide dose response analysis in H209 cancer cells alone or in co-culture with fibroblasts was assessed using the IncuCyte S3 software. Left panel shows the time course increase in apoptotic cancer cells number identified as Annexin V-positive cells using IncuCyte S3 live microscopy. Graphs are presented as fold increase from T0 using the “Red Area / Green Area Normalized to T0” software analysis. Right panel shows AUC computed from “Fold increase Annexin V-positive cells” *vs* time using the trapezoidal rule across 72 hours. Statistical significance between H209 alone (SCLC alone) and H209 in co-culture (Direct co-culture) groups was assessed using one-way ANOVA with Šidák’s post hoc correction. Data are shown as mean ± SEM from one representative experiment (6 replicates/plate). p-values are indicated in the graphs: ***, p< 000.5; ****, p< 0.0001. **(C)** H209 cancer cells alone or in co-culture with fibroblasts were treated with the indicated doses of etoposide doses for 3 days. The cancer cells in mixed population were separated using FACS sorting after 3 days of treatment. Protein expression of cleaved and uncleaved caspase -3 and caspase-7 in both conditions was determined by immunoblotting. **(D)** H209 cancer cells in single and co-culture conditions were treated with a YAP1-inhibitor, verteporfin, for 24 hours prior to drug etoposide treatment. The cancer cells in mixed population were separated using FACS sorting after 3 days of treatment. Protein expression of cleaved and uncleaved caspase -3 and caspase-7 in verteporfin-treated and untreated conditions was determined by immunoblotting. **(E)** H209 cancer cells were grown in different conditioned media and left untreated or treated with the indicated doses of etoposide doses for 3 days. Protein expression of cleaved and uncleaved caspase -3 was determined by immunoblotting.

### YAP1 is partially responsible for direct interaction-induced chemotherapy resistance

Hippo signaling pathway is a highly conserved kinase cascade that was first discovered in *Drosophila melanogaster*, critical in regulating organ size and tissue homeostasis [49, 50]. YAP1 (Yes-associated protein 1) and TAZ (transcriptional co-activator with PDZ-binding motif) are the major downstream effectors of this signaling pathway [51] and dysregulation of Hippo pathway is associated with various cancers, such as breast, brain, liver, lung, prostate, gastric, pancreatic, and colorectal [51], liver cancer [52], pancreatic cancer [53], colorectal cancer [54], and lung cancer [55]. However, mutations within the pathway, particularly in the core kinase module components, are relatively rare and are typically present in less than 10% of cancer cases [56]. The role of Hippo signaling in EMT and tumorigenesis is primarily mediated by the hyperactivity of its downstream effectors, YAP1/TAZ. Both *in vitro* and *in vivo* studies have demonstrated that, in addition to cell proliferation, migration, and invasion, overexpression of YAP1 is also associated with chemotherapy resistance. Studies have shown that YAP1 is significantly overexpressed in chemo-resistant esophageal cancer tissues compared to sensitive esophageal cancer tissues. Furthermore, inhibition of YAP1 sensitizes cells to cytotoxic drugs [57].

In our previous studies, we showed that the endogenous expression of YAP1 is markedly low in SCLC lines (H209 and H69); however, it is highly upregulated when the cancer cells are in direct contact with fibroblasts [44]. To determine whether high expression of YAP1 plays a role in chemotherapy resistance in our model, we inhibited YAP1 activity in cancer cells using the YAP1 inhibitor verteporfin [58]. We cultured the green fluorescent labeled cancer cells alone or in contact with fibroblasts for 24h to allow direct contact as before. Following initial contact, we treated the H209 cells alone or in direct co-culture conditions with either etoposide alone at 12.5 μM or etoposide and verteporfin for an additional 48h. Western blot analysis was performed on FACS-sorted treated cells to assess apoptosis in cancer cells (Figure 2D). As evaluated by the level of uncleaved caspase 3, and consistent with the literature, inhibition of YAP1 sensitizes these cells to etoposide treatment (Figure 2D, Mix Etoposide *vs* Etop+Verte). These results suggest that YAP1 activity is partially responsible for chemotherapy resistance in our model.

### Indirect interaction and paracrine factors are not sufficient to induce chemoresistance

We next tested if the effect on chemoresistance in cancer cells could be attributed to direct interaction with fibroblasts or if the paracrine signaling could elicit a similar outcome. For this, cancer cells alone were cultured in (i) fresh media, (ii) conditioned media from CCD8, or (iii) conditioned media from a 3-day mix of CCD8 and H209 and treated with two different doses of etoposide for 72 hours. Western blot analysis was performed to assess the cancer cell death in each condition using cleaved and uncleaved caspase 3 and caspase 7 as read-out. There were no significant differences in the levels of cleaved and uncleaved caspase 3 (Figure 2E) and caspase 7 (Supplemental Figure 2) across all three experimental conditions, suggesting that paracrine signaling alone is insufficient to induce early chemoresistance in our model. Although we previously observed a slight upregulation of YAP1 in response to paracrine signaling when cancer cells were cultured in indirect contact with fibroblasts [44], this seems insufficient to induce early chemoresistance in our model.

### Three-dimensional (3D) culture model recapitulates chemoresistance phenotype in SCLC

Two-dimensional (2D) co-culture systems are highly reproducible and cost-effective. However, they fail to reproduce the complex tumor structure and kinetics [59]. Our 2D model ensures that almost all cancer cells are in contact with fibroblasts (Figure 3A, 2D model), thereby undergoing reprogramming, whereas this might not be the case *in vivo* due to complex tissue cell interactions. To replicate *in vivo* conditions, we developed 3D spheroids by varying the cancer-to-fibroblast cell ratio (Figure 3A, 3D model). For this, we cultured six distinct spheroid conditions, maintaining a constant number of cancer cells while progressively decreasing the number of fibroblasts (fibroblasts-to-cancer cells ratio varies from 1:1 to 1:16). After allowing the spheroids to form over 3 days in 2% Matrigel, we treated them with etoposide at 12.5 μM. We used IncuCyte® Annexin V NIR Dye to label apoptotic cells and acquired live-cell images using the IncuCyte S3 microscope as described (Figure 3B). Quantification of red fluorescence intensity (Annexin V positive cells) across the whole brightfield field (total spheroid area) showed an increase in cell death in spheroids with lower fibroblast number (Figure 3C, left panel, etoposide treatment in 1:16 *vs* 1:8 *vs* 1:2 *vs* 1:1 mix). Because fibroblasts don’t undergo apoptosis at 6.25 μM etoposide (Figure 1D), the quantification of red fluorescence intensity likely reflects H209 cancer cell death within the spheroid. These results are consistent with our observations in the 2D model. Importantly, our results suggest that the greater the fibroblasts infiltration, the higher the chemotherapy resistance (Figure 3C, right panel).

**Figure 3.**
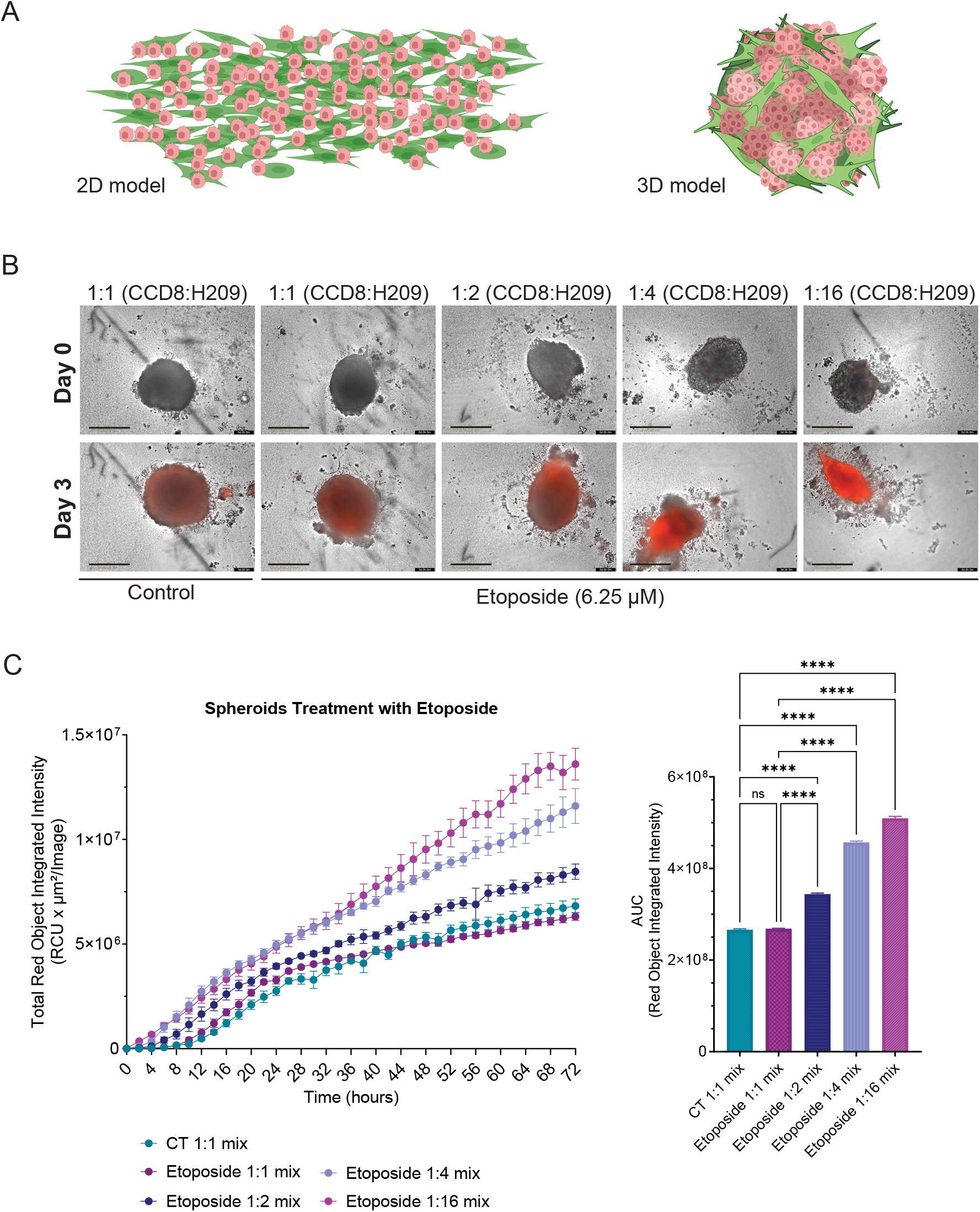
3D model of SCLC cancer cells - lung fibroblasts co-culture. **(A)** The two models of SCLC-fibroblasts co-culture are represented: 2D model-cancer cells and fibroblasts are grown in direct contact with fibroblasts in a 75cm^2^ flask; 3D model: spheroids are created in U-shaped bottom plates by mixing different numbers of cancer cells and fibroblasts to simulate *in vivo* conditions. **(B)** Mix spheroids with different CAFs to cancer cells ratios were treated with the indicated etoposide dose. Annexin V red dye was added at the time of treatment. IncuCyte S3 live microscopy captured the “Brightfield” and “Red fluorescent” spheroid images every 4 hours for 3 days. Microscopy graphs show Day 0 and Day 3 post-treatment in spheroids with different ratios of fibroblasts (CCD8) to cancer cells (H209). Bar = 400 µm. **(C)** Etoposide treatment analysis in H209 cancer cells in mixed spheroids with fibroblasts was assessed using the IncuCyte S3 software. Left panel show the time course increase in the intensity of Annexin V-positive spheroids using IncuCyte S3 live microscopy. Graphs are presented as a time course of “Total Red Object Integrated Intensity (RCU x µm^2^/image)” for 72 hours. Right panel shows AUC computed from “Total Red Object Integrated Intensity” curves *vs* time using the trapezoidal rule across 72 hours. Statistical significance between different spheroid mixed groups was assessed using one-way ANOVA with Šidák’s post hoc correction. Data are shown as mean ± SEM from one representative experiment (6 replicates/plate). p-values are indicated in the graphs: ns, not significant; ****, p< 0.0001.

Altogether, these results suggest that chemoresistance in tumors could begin to develop in cancer cells directly in contact with fibroblasts in the microenvironment, and that the number of infiltrating fibroblasts might be responsible for early and higher chemoresistance.

### *In vitro* drug library screening to uncover new SCLC therapy

We observed that the spheroids composed of 1:1 ratio of fibroblasts to cancer cells exhibited an almost complete resistance to etoposide treatment (Figure 3C). Based on these findings, we selected the 1:1 cancer cells-to-fibroblasts spheroid condition to perform a drug screen using the NIH Drug Clinical Collection to identify potential new therapy that induce apoptosis in mixed spheroids. Spheroids of cancer cells and fibroblasts at a 1:1 ratio were maintained for 3 days to allow spheroid formation. Further, the spheroids were treated with the NIH Drug Clinical collection at a final concentration of 10 μM. Following treatment, spheroids were cultured for an additional 6 to 10 days and monitored using the IncuCyte S3 live-cell imaging system, with images acquired every four hours. Six days post-treatment, the IncuCyte S3 software was used to generate a mask identifying the size of the spheroid and applied across all spheroids within each plate to evaluate the spheroid growth (Figure 4A, yellow mask). Fold growth was calculated as the ratio of spheroid size at day 6 *vs* day 0 (the beginning of treatment) (fold growth= size of spheroid at day 6/ size at day 0).

**Figure 4.**
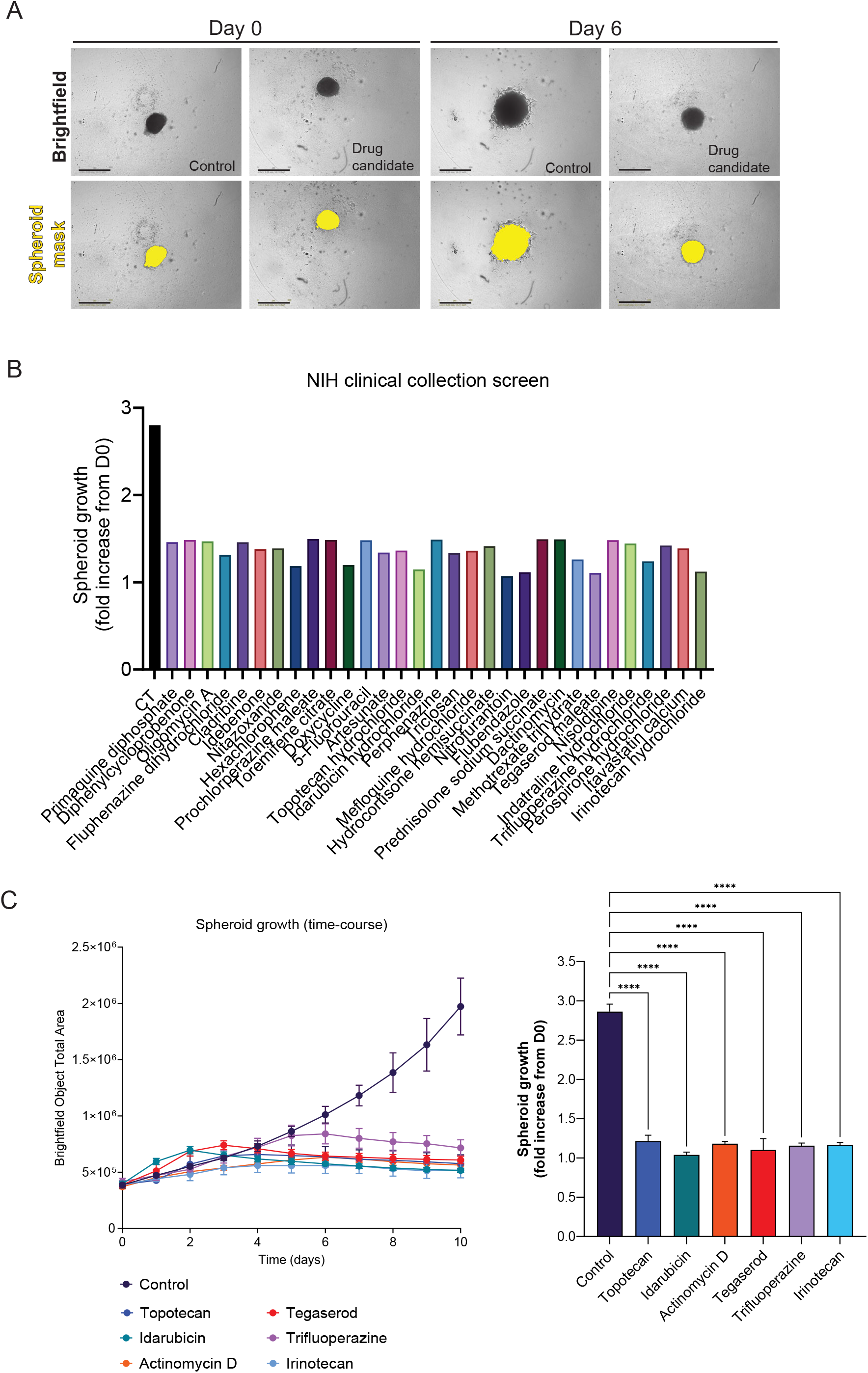
Identification of new therapeutic drugs using high-throughput screening. **(A)** Representative images of “Control” and “Drug candidate” spheroids identified in a high-throughput screening of NIH Clinical Collection. Mixed spheroids of 1:1 ratio of H209 cancer cells and CCD8 lung fibroblasts were established for 3 days before treatment. Spheroid images were captured from Day 0 to Day 6 every 4 hours using IncuCyte S3 live microscopy. At the end of acquisition, a mask (in yellow, Spheroid mask) was designed using the IncuCyte S3 software to determine the size of the spheroid. Bar = 800 µm. **(B)** “Spheroid growth (size fold increase from Day 0)” graph shows all the drugs identified in the first screening as potential growth inhibitors of mixed H209:CCD8 spheroids. A <1.5-fold increase threshold was applied across all the drugs tested from the NIH Clinical Collection. **(C)** Mixed spheroid growth over 10 days for the six drugs with the strongest effect on growth in secondary screenings are presented as a time-course for “Brightfield Object Total Area” as determined by the IncuCyte S3 software (left panel) and “Fold increase from D0” (right panel). Statistical significance between Control and drug treated groups was assessed using one-way ANOVA with Dunnett’s post hoc correction. p-values are indicated in the graphs: ****, p< 0.0001.

Using a 1.5-fold increase as the threshold, we determined that 31 of 707 screened compounds reduced growth to ≤ 1.5-fold in mixed spheroids, whereas the average growth in control wells was 2.8-to 3-fold (Figure 4B). This suggests that the 31 compounds could inhibit growth by either blocking cell replication or inducing apoptosis in cells within the mixed spheroids. We further performed secondary drug screening on the 31 compounds that reduced growth to ≤ 1.5-fold in mixed spheroids, using the same analysis described above. From the secondary screening, we identified six drugs (topotecan, idarubicin, actinomycin D, tegaserod, trifluoperazine, and irinotecan) that suppressed spheroid growth below 1.2-fold over 10 days, indicating near-complete growth inhibition (Figure 4C).

### Idarubicin induces apoptosis in SCLC cells cultured in direct contact with fibroblasts

To determine whether newly identified drugs induce apoptosis in cancer cells or fibroblasts or both, we treated each cell type independently with the selected compounds for 36 hours, and Western Blot analysis was performed to assess apoptosis, as indicated by the presence of cleaved caspase 7 (Figure 5A and 5B). Western blot analysis further revealed that idarubicin, tegaserod, and actinomycin D exerted the strongest pro-apoptotic effects in cancer cells, even exceeding those of the standard-of-care drugs (Figure 5A, H209). We also observed that most of the identified drugs selectively induced apoptosis in cancer cells but not in fibroblasts. Idarubicin, however, also affected fibroblast viability (Figure 5B, CCD8), suggesting that this agent could be used in cancer therapy to induce cell death in both cancer cells and CAFs in the tumor microenvironment. We therefore prioritized this compound for further investigation.

**Figure 5.**
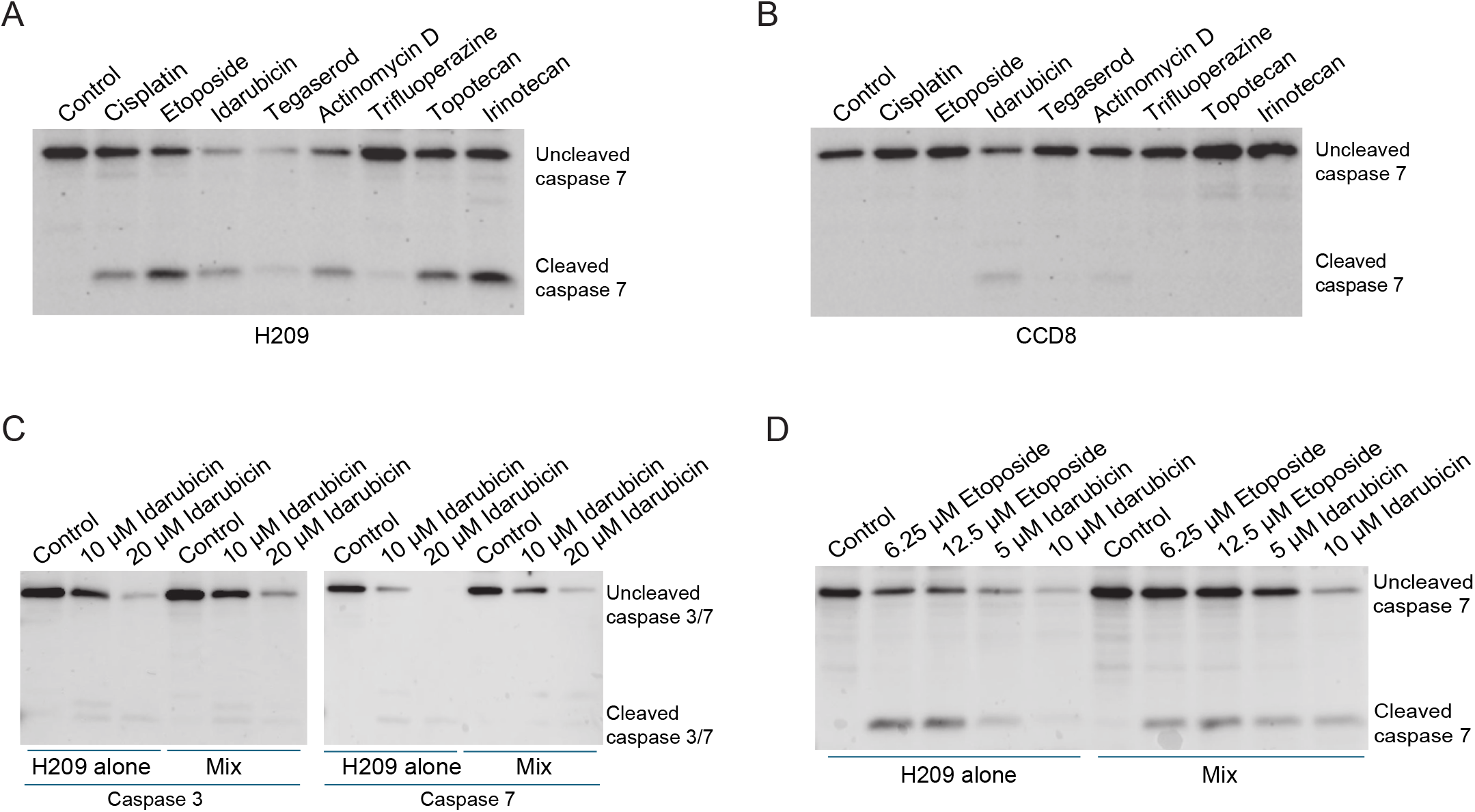
Idarubicin induces apoptosis in H209 cancer cells despite the direct interaction with fibroblasts. **(A-B)** Immunoblot detection of cleaved and uncleaved caspase-3 or caspase-7 in mono-culture of H209 cancer cells **(A)** or CCD8 fibroblasts **(B)** treated with 10µM of drugs identified from drug screening for 36 hours. **(C)** H209 cancer cells alone or in co-culture with fibroblast were treated with idarubicin at two different concentrations and apoptosis in different conditions was determined by immunoblotting detection of cleaved and uncleaved caspase-3 or caspase-7 as described. **(D)** H209 cancer cells alone or in co-culture with fibroblasts were treated with two different doses of etoposide or idarubicin and apoptosis was determined by immunoblotting detection of cleaved and uncleaved caspase-3 or caspase-7.

Next, treatment of H209 cancer cells, either alone or in direct contact with CCD8 fibroblasts, with idarubicin indicated that idarubicin strongly affected cancer cells despite interactions with fibroblasts, particularly at higher doses (Figure 5C). We further compare the response to etoposide and idarubicin at lower doses. We treated cancer cells, either alone or in contact with fibroblasts, with idarubicin at 5 and 10 μM or etoposide at 6.25 and 12.5 μM, as before, for 24 hours. Post-treatment and after FACS sorting, apoptosis was assessed by Western blot using cleaved and uncleaved caspase 7 (Figure 5D). The results showed that cancer cells treated with lower doses of idarubicin are undergoing apoptosis even when in direct contact with fibroblasts (Figure 5D, 5μm and 10μM idarubicin in H209 alone *vs* Mix), while the etoposide-treated cells are more resistant as previously demonstrated. These findings were further validated by an apoptosis assay using flow cytometry (Figure 6 and Supplemental Figure 3). Analysis of early and late apoptosis showed that while direct contact with fibroblasts drastically reduces the percentages of early and late apoptosis in H209 cells after etoposide treatment (Figure 6A-B, etoposide at 6.25 and 12.5 µM in H209 alone *vs* H209 mix), idarubicin induces strong apoptosis under all conditions (Figure 6A-B, idarubicin at 5 and 10 µM in H209 alone *vs* H209 mix). In addition, idarubicin induces strong apoptosis in CCD8 fibroblasts as well (Figure 6C). This data suggests that idarubicin could be an alternative treatment for SCLCs that have undergone EMT and may exhibit chemoresistance when in direct contact with fibroblasts.

**Figure 6.**
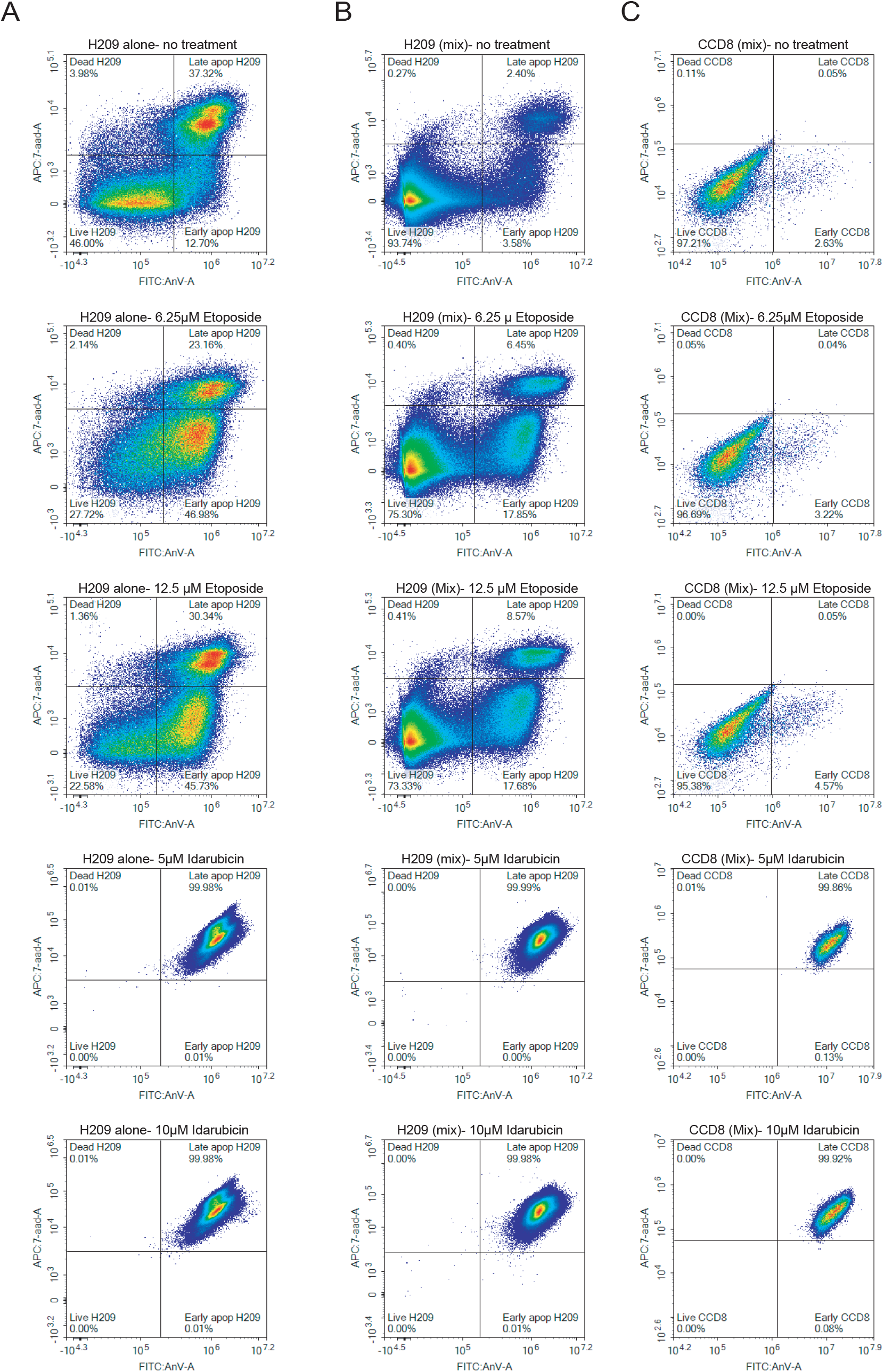
Flow cytometric analysis of apoptosis in mono-culture and co-culture of cancer cells and fibroblasts. Cancer cells cultured alone or in co-culture with fibroblasts were treated with different doses of etoposide and idarubicin as indicated. Apoptosis was assessed using the FITC Annexin V Apoptosis Detection Kit with 7-AAD in H209 cancer cells alone **(A)**, H209 cancer cells from co-culture **(B)**, and CCD8 fibroblasts from co-culture **(C)**. Quadrant gating distinguishes cell populations as live (Annexin V^−^/7-AAD^−^), early apoptotic (Annexin V^+^/7-AAD^−^), late apoptotic (Annexin V^+^/7-AAD^+^), or dead (Annexin V^−^/7-AAD^+^).

## Discussion

Cancer-associated fibroblasts (CAFs) play a critical role in inducing EMT, metastasis, and overall cancer progression. We have previously shown that the interaction between cancer cells and fibroblasts, specifically direct physical interaction, leads to the upregulation of EMT transcription factors such as ZEB1, ZEB2, and TWIST2 in cancer cells [44, 45]. The direct interaction also leads to the upregulation of YAP1, a major regulator of EMT and activator of mesenchymal genes, whose role has been well established in inducing chemoresistance in various cancer types [60-62]. Additionally, we have shown that this direct interaction seems to be required to initiate the paracrine signaling, suggesting that the physical contact between cancer cells and fibroblasts may be playing an early role in EMT, with paracrine signals functioning as a secondary response mechanism [44]. Here, we have extended our investigation to demonstrate that the direct interaction between cancer cells and fibroblasts induces early chemoresistance in cancer cells.

We established a model in which SCLC suspension cells are grown alone or in direct contact with adherent lung fibroblasts. As our previous model showed, SCLC lines adhere to the fibroblasts within a few hours post direct co-culture [44]. Using cisplatin and etoposide as the standard-of-care drugs in SCLC, we first determined the optimal drug doses (12.5 μM and 6.25 μM) that induce apoptosis in cancer cells but not in fibroblasts. Treating cancer cells alone or in mixed direct co-culture with fibroblasts at optimal drug doses revealed that cancer cells in direct contact with fibroblasts are resistant to chemotherapeutic agents, as indicated by decreased numbers of apoptotic cells (Figure 2B and C). This acquired resistance can be attributed to direct interaction between SCLC cells and lung fibroblasts, as cancer cells grown in conditioned media from CCD8 alone or from mixed co-cultures of H209 and CCD8 did not develop early drug resistance, suggesting that paracrine signals alone are insufficient to induce chemoresistance (Figure 2E). Interestingly, when YAP1 activity was inhibited with verteporfin, cancer cells were less resistant to etoposide, indicating that therapy resistance during direct contact with fibroblasts is partially driven by YAP1, and that inhibition of YAP1 resensitizes cells to chemotherapeutic agents (Figure 2D).

We further advanced our study by transitioning from a 2D co-culture system to a more physiologically relevant 3D spheroid model with varying numbers of infiltrating fibroblasts within cancer cell spheroids, revealing that spheroids with a higher number of infiltrating fibroblasts are more resistant to treatment (Figure 3B and 3C). These findings support our hypothesis and is consistent with the previous studies that fibroblasts play a protective role against the chemotherapeutic agents [63-65]. Using the 3D spheroid model, we performed a high-throughput drug screening using NIH Clinical Collection Drugs, with a total of 707 drugs, and identified idarubicin as a therapeutic candidate that induces cell death in cancer cells even when they are in direct contact with the fibroblasts (Figures 5 and 6).

Our findings are significant for several conceptual and translational reasons. While the roles of CAFs-derived exosomes in chemoresistance have been well reported in many studies [21, 66-69], how direct physical interaction between CAFs and cancer cells induces chemoresistance remains poorly understood. Beyond confirming earlier reports on CAF-driven chemoresistance, our study advances the field by offering new evidence that strengthens and expands upon these observations. First, our study demonstrates that physical interaction between CAFs and cancer cells induces early chemoresistance. While we observed some EMT reprogramming in cancer cells when they were grown in conditioned media from the mixed co-culture of cancer cells and fibroblasts in our previous study [44], chemoresistance was not induced in cancer cells under the same conditions (Figure 2E). This suggests that extensive EMT reprogramming, such as that induced by direct interaction [44], is required to confer therapeutic resistance. This finding reveals a novel therapeutic avenue that targets direct cell-cell interactions rather than solely targeting secreted factors. By intervening at the level of physical cellular crosstalk, this approach may help overcome acquired chemoresistance, one of the most significant challenges in cancer progression.

Our study indicates that the observed therapeutic resistance is partially induced through the upregulation of YAP1, which is typically expressed at low levels in SCLC but becomes highly upregulated when cancer cells are in direct contact with fibroblasts [44]. Inhibition of YAP1 restores the sensitivity of cancer cells to therapeutic agents, consistent with previous studies [61], demonstrating a similar effect across several cancers, including triple-negative breast cancer [70], lung cancer [71], and urothelial cancer [72]. Studies have shown that currently available YAP1-inhibitory agents, such as verteporfin, dasatinib, evodiamine, and statins, can be used in combination with anticancer drugs to improve cancer therapeutic efficacy [61]. Upregulation of YAP1 is a common occurrence across many types of cancer and is almost invariably associated with the development of chemoresistance. Our study demonstrates that YAP1 upregulation is driven by the direct interaction between cancer cells and fibroblasts. Thus, targeting the cell-cell crosstalk between the two cell lines may provide a promising therapeutic approach to prevent the YAP1-mediated chemoresistance and enhance the efficacy of chemotherapy.

SCLC is a highly aggressive malignancy that initially responds well to therapy but almost consistently develops chemoresistance [73, 74]. Our results indicate that while direct contact between SCLC and fibroblasts induces early chemoresistance to standard-of-care drugs, this is not the case for idarubicin treatment. This is a critical finding because TME-driven chemoresistance is a major cause of treatment failure in cancer patients in general and SCLC as well. By showing that the idarubicin bypasses early resistance induced by direct cancer cell-fibroblast interactions and remains effective despite fibroblast-mediated protection, our study identifies a therapeutic vulnerability that may improve treatment efficacy in stroma-rich tumors, potentially reduce relapse, and position this drug as a promising therapeutic approach in combination with the standard-of-care. Although both etoposide and idarubicin target Topoisomerase II, their distinct mechanisms of action may underlie the different sensitivity observed in our model. Etoposide induces replication-associated DNA damage by stabilizing Topoisomerase II cleavage complexes, a process that may be attenuated in cancer cells in direct contact with fibroblasts exhibiting altered cell-cycle dynamics and survival signaling. In contrast, idarubicin combines Topoisomerase II inhibition with DNA intercalation and oxidative stress, potentially bypassing the protective adaptations and maintaining cytotoxic efficacy despite fibroblast-mediated resistance. One caveat of our study is that, although idarubicin appears to be more effective than current standard-of-care drugs at inducing SCLC apoptosis despite direct contact with fibroblasts, we have not tested it under conditions in which cancer cells are in long-term contact with fibroblasts. A phase 2 study previously reported that oral idarubicin showed limited efficacy as a first-line treatment for extensive-stage small cell lung cancer [75]. The study also highlighted concerns regarding the appropriateness of testing single-agent therapies in a highly aggressive disease where combination chemotherapy was already showing superior response rates. However, with advances in drug delivery systems, pharmacokinetics, and patient-personalized strategies, idarubicin could potentially be evaluated more rigorously today, particularly in tumors that have developed greater self-sufficiency and acquired stromal-like functions during disease progression or relapse through interactions with fibroblasts in the TME.

## Conclusions

Overall, our findings shift the therapeutic perspective from targeting primarily exosomes and secreted factors to interfering with cellular interactions between cancer cells and components of the tumor microenvironment, which appear to be upstream regulatory mechanisms driving the signaling pathways that induce chemoresistance. In this context, direct physical contact between cancer cells and fibroblasts emerges as a critical initiating event that promotes transcriptional reprogramming, including YAP1 activation and EMT-associated changes, ultimately enabling early adaptation to therapy. Perhaps cancer cells that come into early contact with the stroma are the first to develop resistance and remain resistant for the duration of treatment. Our 3D model of interaction could be used as a strategy to screen additional drug libraries with the potential to uncover novel therapies that prevent or delay the onset of microenvironment-driven drug resistance.

## Materials and methods

### Cell culture conditions

Small cell lung carcinoma cell lines H209 (HTB-172) and H69 (HTB-119), and normal lung fibroblast CCD8 (CCL-201), were obtained from ATCC and cultured in RPMI medium supplemented with 10% FBS. Cells were cultured at 37° C in a 5% CO2 incubator. All cell lines were authenticated by short tandem repeat (STR) profiling. All cell lines were tested for *Mycoplasma* contamination. Two types of co-cultures were performed. For monoculture conditions, the cells were grown alone. For the 2D co-culture condition or “Mix”, fibroblasts and cancer cells were cultured in the same flask in a fibroblasts: cancer cells ratio of 1:6 unless otherwise noted. The fibroblasts were seeded 24 hours prior to the addition of H209 and H69 cells to allow them to adhere. When indicated, Verteporfin (Cayman Chemical #17334) was used at a concentration of 500 nM prior to chemotherapy treatment. H209 cancer cells and CCD8 fibroblasts were labeled with either red (PKH26 Red Fluorescent Linker Kit) or green (PKH67 Green Fluorescent Linker Kit) from Sigma according to the manufacturer’s instructions prior to co-culture and treatment with various drugs and FACS sorting.

### Drug treatment and high-throughput microscopy analysis

Standard-of-care drugs for SCLC, cisplatin and etoposide, were used at different concentrations to determine the optimal doses that induce cell death in cancer cells but not fibroblasts. H209, H69, and CCD8 cells alone were treated with serial dilutions (50μM, 25 μM, 12.5 μM, 6.25 μM, and 3.125 μM) of cisplatin alone and etoposide alone. For the live apoptosis assay, only cancer cells were labeled using the PKH67 Green Fluorescent Linker (Sigma). Apoptotic cells were labeled using the IncuCyte® Annexin V NIR Dye (Annexin Red), and live cell images were acquired using the IncuCyte S3 microscope every 4 hours. Drug dose-response was evaluated using the integrated software of the IncuCyte S3 microscope. Red Area / Phase Area Normalized to T0 metric was used to measure the increase in apoptotic cells in single cultures and the Red Area / Green Area Normalized to T0 metric was used to measure the increase in apoptotic cancer cells in mixed cultures.

### Conditioned media

Conditioned media from either the monoculture of CCD8 or the co-culture of CCD8 and H209, cultured for 72 hours, were collected. Cancer cells grown in either fresh media, conditioned media from CCD8, or conditioned media from the co-culture were treated with optimum doses of etoposide for 72 hours.

### Culture of 3D spheroid model

Six distinct spheroid model conditions were established, maintaining a constant number of cancer cells while progressively decreasing the number of fibroblasts. H209 cancer cells and CCD8 fibroblasts were mixed at fibroblasts-to-cancer cells ratios ranging from 1:1 to 1:16 in 2% Matrigel in a 96-well round-bottom ultra-low attachment (ULA) plates (Corning #7007). The mixture was cultured for 3 days to allow spheroid formation prior to drug treatment.

### Western Blot analysis

Protein extraction was performed using a RIPA lysis buffer. Bicinchoninic Acid (BCA) Assay was employed to measure protein concentration. SDS-PAGE was performed on 15% gradient gels to separate proteins by molecular weight, using the same protein concentration across all conditions. After the electrophoresis, the gel was transferred to a nitrocellulose membrane, and the membranes were blocked with 5% milk in Tris-buffered saline with Tween-20 (TBS-T) for 60 minutes at room temperature prior to incubation with primary antibody overnight at 4°C. Following primary antibody the membranes were incubated with the secondary antibody for 1 hour at room temperature. The primary antibodies used were caspase 3 (cat #9662) and caspase 7 (cat #9492) from Cell Signaling. Secondary antibodies horseradish peroxidase (HRP)-anti-mouse and anti-rabbit (1:5,000) were from Cell Signaling.

### Cell labeling and FACS sorting

H209 or H69 cancer cells and CCD8 fibroblasts were labeled with PKH26 Red Fluorescent Linker Kit or the PKH67 Green Fluorescent Linker Kit (Millipore Sigma), per manufacturer’s instructions. Briefly, the cells were collected, washed twice in serum-free medium, and resuspended in 1mL PKH diluent. The PKH26 and PKH67 dyes were diluted separately in PKH diluent (4μL/1mL diluent). Cells and dye were mixed and incubated for 8 min at room temperature. Staining was stopped by adding 10 mL of RPMI supplemented with 10% FBS. Cells were washed 2 more times with complete media before plating in monoculture or co-culture. Following co-culture and treatment with various drugs at different time points, mixed cells were detached with Trypsin, washed, and sorted by fluorescent marker using the Bigfoot Spectral Cell Sorter (Invitrogen).

### Apoptosis assay

For apoptosis detection by FACS cells were treated with the corresponding chemotherapy for the duration indicated and apoptosis was determined with the FITC Annexin V Apoptosis Detection Kit with 7-AAD (BioLegend # 640922), per manufacturer’s instructions. Briefly, the cells were washed twice with cold Cell Staining Buffer and then resuspend it in Annexin V Binding Buffer at a concentration of 0.25-1.0 x 10^7^ cells/mL. Approximately 100 µL of cell suspension were transfer in 5 mL test tube and 5 µL of fluorochrome conjugated Annexin V was added together with 7-AAD viability dye. Cells were incubate for 15 min at RT (25°C) in the dark. Annexin V Binding Buffer was added to each tube and samples were analyzed by flow cytometry.

### Drug library high-throughput screening

H209 cancer cells and CCD8 fibroblasts were mixed in a 1:1 ratio (10,000 cells/well) in 2% Matrigel in a 96-well round-bottom ULA plate. The cells were cultured for 72 hours to allow spheroid formation. The established spheroids were then treated with the NIH Clinical Collection at a final concentration of 10 μM. The drug-treated spheroid plates were placed in the Incucyte S3 microscope, and images were acquired every 4 hours for 6 to 10 days. Incucyte S3 software was used to design a mask to determine the area of the brightfield object (single spheroid). For subsequent validation all drugs were purchased from Selleck Chemicals and used as indicated.

### Statistical analysis

All Western blot experiments were performed at least three times. High-throughput drug screening was performed once to identify the target drugs. Following initial screening, the “daughter plates” with the target drugs were used for validation. All Incucyte S3 live acquisition experiments were performed at least twice, with 6 replicates per condition on each plate. Data are presented as representative curves and area under the curve (AUC) graphs. For AUC significance, multiparametric ANOVA analysis was performed as indicated using Prism v9 (GraphPad Software). Values of *P* < 0.005 were considered significant, values of *P* < 0.001 were considered highly significant.

## Supporting information

Original Western blots

## Author Contributions

AMD and ND conceptualized the project and acquired funds for the project. ND, BL, DC and AMD performed experimental work and data analysis. AMD and ND wrote the original draft, reviewed and edited the final manuscript.

## Acknowledgments

We would like to acknowledge the services offered by the Center of Applied Immunology and Pathological Processes Immunophenotyping Core (CAIPP) (RRID: SCR_024781) and are grateful to Dr. Sushma Bharrhan for providing reagents and technical assistance with apoptosis FACS analysis. Live imaging experiments were performed in the INLET Core at the LSU Health Sciences Center-Shreveport and Feist-Weiller Cancer Center (RRID:SCR_024990). FASC sorting and analysis were performed in the Research Core Facility at LSU Health Sciences Center-Shreveport (RRID:SCR_024775).

## References

1. Dhiman, V.K., M. Kumari, and D. Singh, Chemoresistance: The hidden barrier in cancer treatment. Cancer Pathog Ther, 2026. 4(2): p. 98–109.

2. Bray, F., et al., Global cancer statistics 2022: GLOBOCAN estimates of incidence and mortality worldwide for 36 cancers in 185 countries. CA Cancer J Clin, 2024. 74(3): p. 229–263.

3. Ramos, A., S. Sadeghi, and H. Tabatabaeian, Battling Chemoresistance in Cancer: Root Causes and Strategies to Uproot Them. Int J Mol Sci, 2021. 22(17).

4. Yeldag, G., A. Rice, and A. Del Río Hernández, Chemoresistance and the Self-Maintaining Tumor Microenvironment. Cancers (Basel), 2018. 10(12).

5. Liu, B., et al., Protecting the normal in order to better kill the cancer. Cancer Med, 2015. 4(9): p. 1394–403.

6. Zheng, H.C., The molecular mechanisms of chemoresistance in cancers. Oncotarget, 2017. 8(35): p. 59950–59964.

7. Brasseur, K., N. Gévry, and E. Asselin, Chemoresistance and targeted therapies in ovarian and endometrial cancers. Oncotarget, 2017. 8(3): p. 4008–4042.

8. Longley, D.B. and P.G. Johnston, Molecular mechanisms of drug resistance. J Pathol, 2005. 205(2): p. 275–92.

9. Kannampuzha, S. and A.V. Gopalakrishnan, Cancer chemoresistance and its mechanisms: Associated molecular factors and its regulatory role. Med Oncol, 2023. 40(9): p. 264.

10. Krivak, T.C., et al., Relationship between ERCC1 polymorphisms, disease progression, and survival in the Gynecologic Oncology Group Phase III Trial of intraperitoneal versus intravenous cisplatin and paclitaxel for stage III epithelial ovarian cancer. J Clin Oncol, 2008. 26(21): p. 3598–606.

11. Oliver, T.G., et al., Chronic cisplatin treatment promotes enhanced damage repair and tumor progression in a mouse model of lung cancer. Genes Dev, 2010. 24(8): p. 837–52.

12. Dogan, E., et al., Targeting Apoptosis to Overcome Chemotherapy Resistance, in Metastasis, C.M. Sergi, Editor. 2022, Exon Publications Copyright: The Authors.; The authors confirm that the materials included in this chapter do not violate copyright laws. Where relevant, appropriate permissions have been obtained from the original copyright holder(s), and all original sources have been appropriately acknowledged or referenced.: Brisbane (AU).

13. Haq, R. and B. Zanke, Inhibition of apoptotic signaling pathways in cancer cells as a mechanism of chemotherapy resistance. Cancer Metastasis Rev, 1998. 17(2): p. 233–9.

14. Fernald, K. and M. Kurokawa, Evading apoptosis in cancer. Trends Cell Biol, 2013. 23(12): p. 620–33.

15. Khan, S.U., et al., Unveiling the mechanisms and challenges of cancer drug resistance. Cell Commun Signal, 2024. 22(1): p. 109.

16. Mansoori, B., et al., The Different Mechanisms of Cancer Drug Resistance: A Brief Review. Adv Pharm Bull, 2017. 7(3): p. 339–348.

17. Jena, B.C., et al., Cancer associated fibroblast mediated chemoresistance: A paradigm shift in understanding the mechanism of tumor progression. Biochim Biophys Acta Rev Cancer, 2020. 1874(2): p. 188416.

18. Teicher, B.A., et al., Tumor resistance to alkylating agents conferred by mechanisms operative only in vivo. Science, 1990. 247(4949 Pt 1): p. 1457–61.

19. McMillin, D.W., J.M. Negri, and C.S. Mitsiades, The role of tumour-stromal interactions in modifying drug response: challenges and opportunities. Nat Rev Drug Discov, 2013. 12(3): p. 217–28.

20. Meads, M.B., R.A. Gatenby, and W.S. Dalton, Environment-mediated drug resistance: a major contributor to minimal residual disease. Nat Rev Cancer, 2009. 9(9): p. 665–74.

21. Zhang, H., et al., CAF secreted miR-522 suppresses ferroptosis and promotes acquired chemo-resistance in gastric cancer. Mol Cancer, 2020. 19(1): p. 43.

22. Ren, J., et al., Carcinoma-associated fibroblasts promote the stemness and chemoresistance of colorectal cancer by transferring exosomal lncRNA H19. Theranostics, 2018. 8(14): p. 3932–3948.

23. Qu, Z., et al., CAFs-secreted exosomal cricN4BP2L2 promoted colorectal cancer stemness and chemoresistance by interacting with EIF4A3. Exp Cell Res, 2022. 418(2): p. 113266.

24. Qi, R., et al., Cancer-associated fibroblasts suppress ferroptosis and induce gemcitabine resistance in pancreatic cancer cells by secreting exosome-derived ACSL4-targeting miRNAs. Drug Resist Updat, 2023. 68: p. 100960.

25. Hu, J.L., et al., CAFs secreted exosomes promote metastasis and chemotherapy resistance by enhancing cell stemness and epithelial-mesenchymal transition in colorectal cancer. Molecular Cancer, 2019. 18(1): p. 91.

26. Hashemi, M., et al., EMT mechanism in breast cancer metastasis and drug resistance: Revisiting molecular interactions and biological functions. Biomed Pharmacother, 2022. 155: p. 113774.

27. Sommers, C.L., et al., Loss of epithelial markers and acquisition of vimentin expression in adriamycin- and vinblastine-resistant human breast cancer cell lines. Cancer Res, 1992. 52(19): p. 5190–7.

28. Du, B. and J.S. Shim, Targeting Epithelial-Mesenchymal Transition (EMT) to Overcome Drug Resistance in Cancer. Molecules, 2016. 21(7).

29. Ebrahimi, N., et al., Harnessing function of EMT in cancer drug resistance: a metastasis regulator determines chemotherapy response. Cancer Metastasis Rev, 2024. 43(1): p. 457–479.

30. Teicher, B.A., et al., Transforming growth factor-beta in in vivo resistance. Cancer Chemother Pharmacol, 1996. 37(6): p. 601–9.

31. Wu, Y., et al., Expression of Wnt3 activates Wnt/β-catenin pathway and promotes EMT-like phenotype in trastuzumab-resistant HER2-overexpressing breast cancer cells. Mol Cancer Res, 2012. 10(12): p. 1597–606.

32. Saxena, M., et al., Transcription factors that mediate epithelial-mesenchymal transition lead to multidrug resistance by upregulating ABC transporters. Cell Death Dis, 2011. 2(7): p. e179.

33. Siebzehnrubl, F.A., et al., The ZEB1 pathway links glioblastoma initiation, invasion and chemoresistance. EMBO Mol Med, 2013. 5(8): p. 1196–212.

34. Haslehurst, A.M., et al., EMT transcription factors snail and slug directly contribute to cisplatin resistance in ovarian cancer. BMC Cancer, 2012. 12: p. 91.

35. Liu, L., et al., Inhibition of Wnt/β-catenin pathway reverses multi-drug resistance and EMT in Oct4(+)/Nanog(+) NSCLC cells. Biomed Pharmacother, 2020. 127: p. 110225.

36. Jackman, D.M. and B.E. Johnson, Small-cell lung cancer. Lancet, 2005. 366(9494): p. 1385–96.

37. Rudin, C.M., et al., Small-cell lung cancer. Nat Rev Dis Primers, 2021. 7(1): p. 3.

38. Glynn, S.E., et al., Extrapulmonary Small Cell Carcinoma: A Single-Institution Review of Brain Metastases, Treatment Paradigms, and Patient Outcomes. Cureus, 2025. 17(1): p. e76974.

39. Nishino, K., et al., Small-cell lung carcinoma with long-term survival: A case report. Oncol Lett, 2011. 2(5): p. 827–830.

40. Herzog, B.H., S. Devarakonda, and R. Govindan, Overcoming Chemotherapy Resistance in SCLC. J Thorac Oncol, 2021. 16(12): p. 2002–2015.

41. Chen, P., et al., Mechanisms of drugs-resistance in small cell lung cancer: DNA-related, RNA-related, apoptosis-related, drug accumulation and metabolism procedure. Transl Lung Cancer Res, 2020. 9(3): p. 768–786.

42. Fridman, R., et al., Reconstituted basement membrane (matrigel) and laminin can enhance the tumorigenicity and the drug resistance of small cell lung cancer cell lines. Proc Natl Acad Sci U S A, 1990. 87(17): p. 6698–702.

43. Lu, Y., et al., Dynamic phenotypic reprogramming and chemoresistance induced by lung fibroblasts in small cell lung cancer. Sci Rep, 2024. 14(1): p. 2884.

44. Dhungel, N., et al., Assessing the epithelial-to-mesenchymal plasticity in a small cell lung carcinoma (SCLC) and lung fibroblasts co-culture model. Front Mol Biosci, 2023. 10: p. 1096326.

45. Dhungel, N. and A.M. Dragoi, Exploring the multifaceted role of direct interaction between cancer cells and fibroblasts in cancer progression. Front Mol Biosci, 2024. 11: p. 1379971.

46. Stinchcombe, T.E. and E.M. Gore, Limited-stage small cell lung cancer: current chemoradiotherapy treatment paradigms. Oncologist, 2010. 15(2): p. 187–95.

47. Farago, A.F. and F.K. Keane, Current standards for clinical management of small cell lung cancer. Transl Lung Cancer Res, 2018. 7(1): p. 69–79.

48. Behrouzi, R. and F. Blackhall, State of the art in treatment of small cell lung cancer. Ther Adv Med Oncol, 2025. 17: p. 17588359251363518.

49. Zhou, T., et al., The Hippo/YAP signaling pathway: the driver of cancer metastasis. Cancer Biol Med, 2023. 20(7): p. 483–9.

50. Piccolo, S., S. Dupont, and M. Cordenonsi, The biology of YAP/TAZ: hippo signaling and beyond. Physiol Rev, 2014. 94(4): p. 1287–312.

51. Akrida, I., V. Bravou, and H. Papadaki, The deadly cross-talk between Hippo pathway and epithelial-mesenchymal transition (EMT) in cancer. Mol Biol Rep, 2022. 49(10): p. 10065–10076.

52. Wu, H., et al., The role of YAP1 in liver cancer stem cells: proven and potential mechanisms. Biomark Res, 2022. 10(1): p. 42.

53. Liu, M., et al., Zinc-Dependent Regulation of ZEB1 and YAP1 Coactivation Promotes Epithelial-Mesenchymal Transition Plasticity and Metastasis in Pancreatic Cancer. Gastroenterology, 2021. 160(5): p. 1771–1783.e1.

54. Jin, L., et al., YAP inhibits autophagy and promotes progression of colorectal cancer via upregulating Bcl-2 expression. Cell Death Dis, 2021. 12(5): p. 457.

55. Yang, Q., et al., LINC02159 promotes non-small cell lung cancer progression via ALYREF/YAP1 signaling. Mol Cancer, 2023. 22(1): p. 122.

56. Cunningham, R. and C.G. Hansen, The Hippo pathway in cancer: YAP/TAZ and TEAD as therapeutic targets in cancer. Clin Sci (Lond), 2022. 136(3): p. 197–222.

57. Song, S., et al., The Hippo Coactivator YAP1 Mediates EGFR Overexpression and Confers Chemoresistance in Esophageal Cancer. Clin Cancer Res, 2015. 21(11): p. 2580–90.

58. Lui, J.W., et al., The Efficiency of Verteporfin as a Therapeutic Option in Pre-Clinical Models of Melanoma. J Cancer, 2019. 10(1): p. 1–10.

59. Li, X., C. González-Maroto, and M. Tavassoli, Crosstalk between CAFs and tumour cells in head and neck cancer. Cell Death Discov, 2024. 10(1): p. 303.

60. Huang, W., et al., PPM1G promotes chemoresistance in triple negative breast cancer by enhancing YAP signaling. Pharmacol Res, 2025. 221: p. 107978.

61. Shibata, M., K. Ham, and M.O. Hoque, A time for YAP1: Tumorigenesis, immunosuppression and targeted therapy. Int J Cancer, 2018. 143(9): p. 2133–2144.

62. Yang, R., et al., Endothelin-1 Mediates Oxaliplatin Resistance via Activation of YAP Signaling in Colorectal Cancer. Oncol Res, 2025. 33(12): p. 3945–3971.

63. Zhao, Z., et al., Potential mechanisms of cancer-associated fibroblasts in therapeutic resistance. Biomed Pharmacother, 2023. 166: p. 115425.

64. Fang, W.B., M. Yao, and N. Cheng, Priming cancer cells for drug resistance: role of the fibroblast niche. Front Biol (Beijing), 2014. 9(2): p. 114–126.

65. Masuda, H., Cancer-associated fibroblasts in cancer drug resistance and cancer progression: a review. Cell Death Discov, 2025. 11(1): p. 341.

66. Hu, J.L., et al., CAFs secreted exosomes promote metastasis and chemotherapy resistance by enhancing cell stemness and epithelial-mesenchymal transition in colorectal cancer. Mol Cancer, 2019. 18(1): p. 91.

67. Zhuang, J., et al., Cancer-Associated Fibroblast-Derived miR-146a-5p Generates a Niche That Promotes Bladder Cancer Stemness and Chemoresistance. Cancer Res, 2023. 83(10): p. 1611–1627.

68. Liu, M., T.Z. Li, and C. Xu, The role of tumor-associated fibroblast-derived exosomes in chemotherapy resistance of colorectal cancer and its application prospect. Biochim Biophys Acta Gen Subj, 2025. 1869(6): p. 130796.

69. Yang, C., et al., Exosomes derived from cancer-associated fibroblasts promote tumorigenesis, metastasis and chemoresistance of colorectal cancer by upregulating circ_0067557 to target Lin28. BMC Cancer, 2024. 24(1): p. 64.

70. Andrade, D., et al., YAP1 inhibition radiosensitizes triple negative breast cancer cells by targeting the DNA damage response and cell survival pathways. Oncotarget, 2017. 8(58): p. 98495–98508.

71. Cheng, H., et al., Functional genomics screen identifies YAP1 as a key determinant to enhance treatment sensitivity in lung cancer cells. Oncotarget, 2016. 7(20): p. 28976–88.

72. Ooki, A., et al., YAP1 and COX2 Coordinately Regulate Urothelial Cancer Stem-like Cells. Cancer Res, 2018. 78(1): p. 168–181.

73. Zhai, X., et al., Current and future therapies for small cell lung carcinoma. J Hematol Oncol, 2025. 18(1): p. 37.

74. Pietanza, M.C., et al., Small cell lung cancer: will recent progress lead to improved outcomes? Clin Cancer Res, 2015. 21(10): p. 2244–55.

75. Cullen, M.H., et al., Testing new drugs in untreated small cell lung cancer may prejudice the results of standard treatment: a phase II study of oral idarubicin in extensive disease. Cancer Treat Rep, 1987. 71(12): p. 1227–30.

