## Supplementary Figures for "Direct interaction between cancer cells and fibroblasts promotes early chemoresistance to standard-of-care drug therapy in small cell lung cancer (SCLC)"

**Supplemental Figure 1**. **Apoptosis induced by cisplatin and etoposide in H69, H209 cancer cells and CCD8 lung fibroblasts. (A-D)** Graphs show the time course increase in apoptotic cells number identified as Annexin V-positive cells using IncuCyte S3 live microscopy. Graphs are presented as fold increase from T0 using the "Red Area / Phase Area Normalized to T0" software analysis during a 72 hours time course after treatment with cisplatin and etoposide. One representative experiment is shown (6 replicates/plate). **(E)** Immunoblot analysis of cleaved and uncleaved caspase-3 and caspase-7 after cisplatin treatment for 3 days.

**Supplemental Figure 2**. **Apoptosis in H209 cancer cells after treatment with etoposide in conditioned media.** H209 cancer cells were grown in different conditioned media and left untreated or treated with the indicated doses of etoposide doses for 3 days. Protein expression of cleaved and uncleaved caspase -7 was determined by immunoblotting.

**Supplemental Figure 3**. **Gating strategy for FASC analysis of apoptosis. (A)** For analysis of mono-culture of H209 the first gate in the gating strategy is Scatter. From this gate a doublet discrimination gate "Singlets" was created. Finally, from the "Singlets" gate, a bivariate plot with APC:7-aad on the y-axis and FITC:AnV on the x-axis was generated. **(B)** For analysis of co-culture of H209 and fibroblasts the first gate in the gating strategy is a scatter plot with both CCD8 and H209 gated based on size. Next, both populations were gated individually onto doublet discrimination plots labeled CCD8 or H209 singlets. Finally, both singlet gates were plotted onto their respective bivariate plots with APC:7-aad on the y-axis and FITC:AnV on the x-axis.

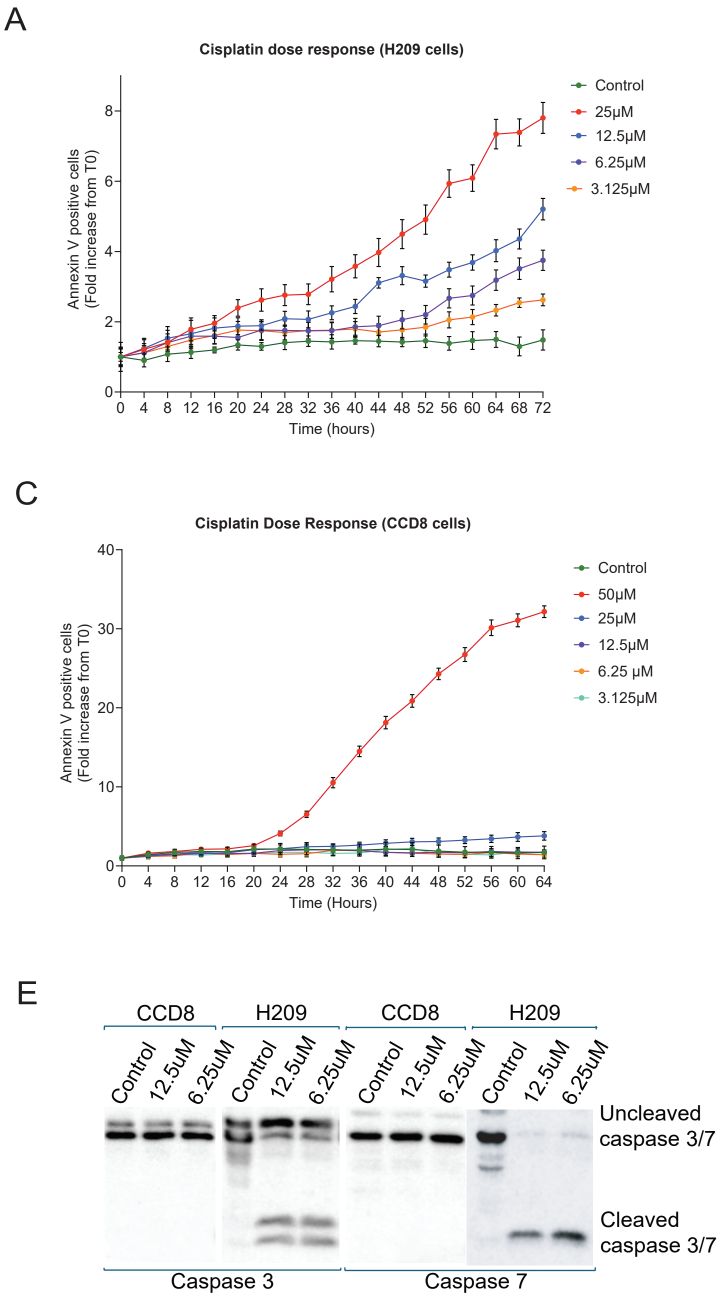


Supplemental Figure 1

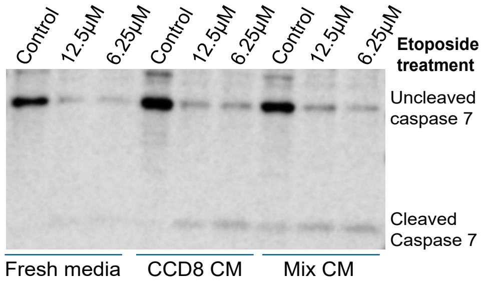


Supplemental Figure 2

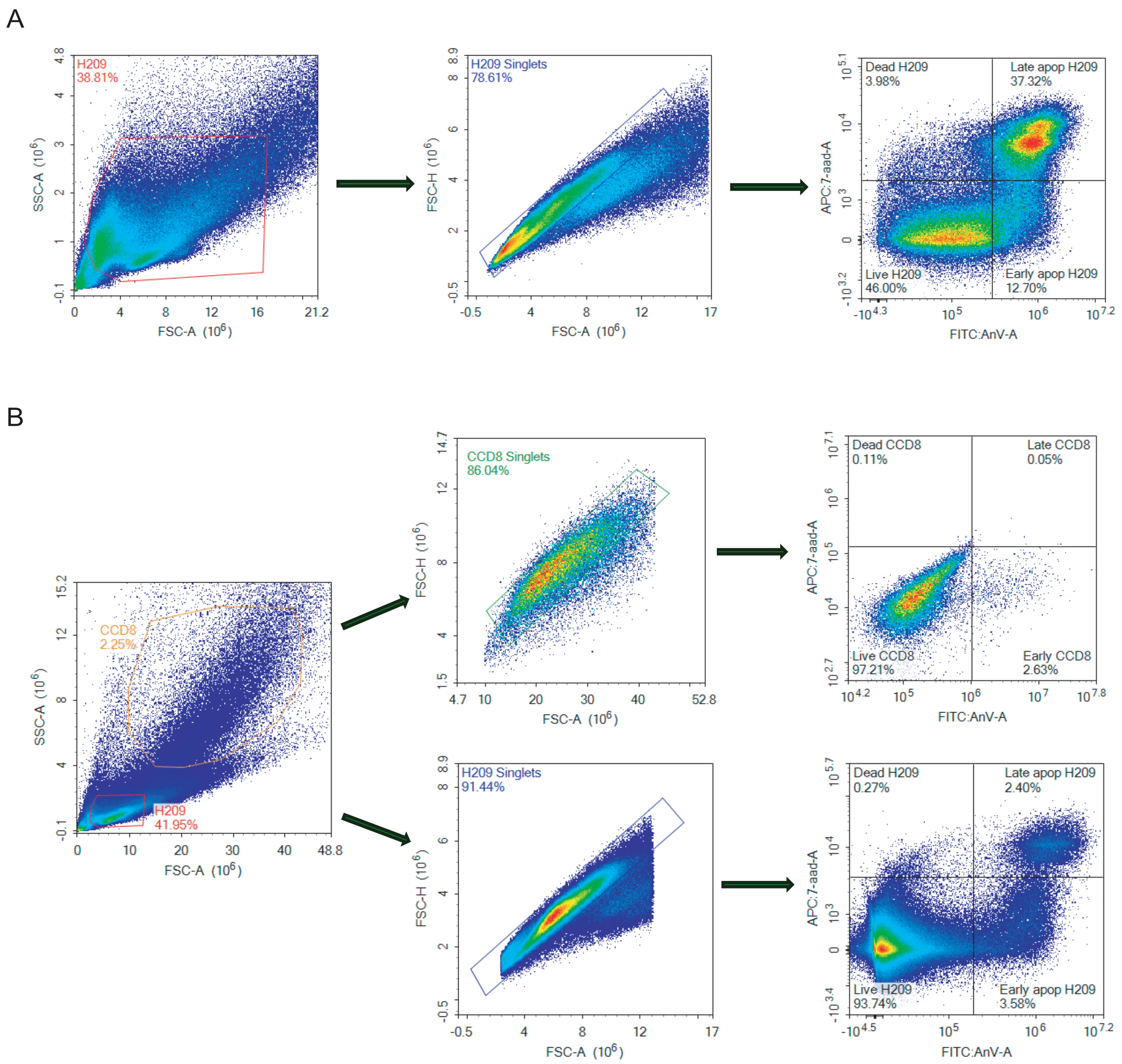


Supplemental Figure 3
