## Supplementary material for "Direct interaction between cancer cells and fibroblasts promotes early chemoresistance to standard-of-care drug therapy in small cell lung cancer (SCLC)": Original Western blots

Figure 1D: Caspase 3 (H69, H209, CCD8 treatment with etoposide)

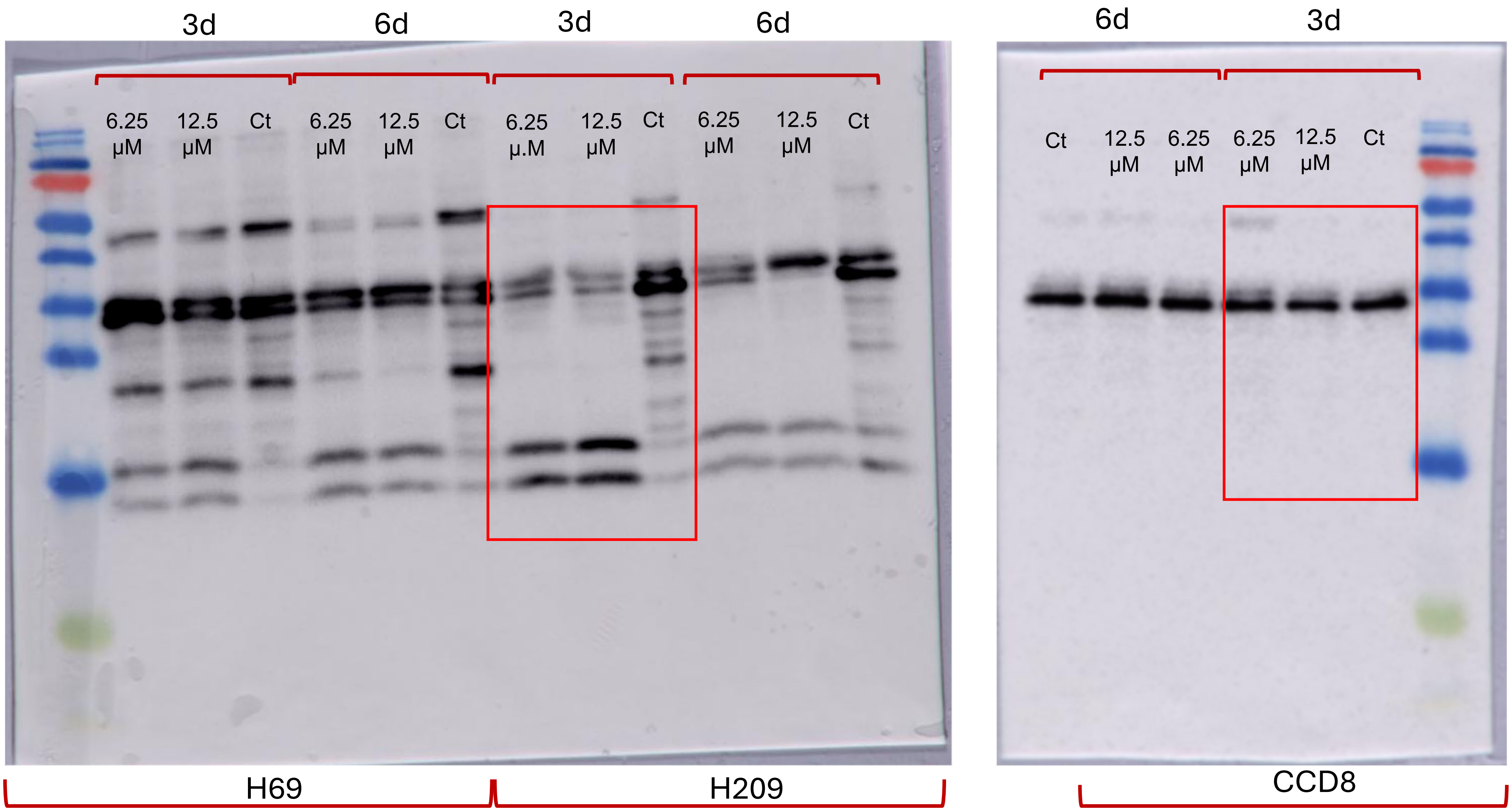

Figure 1D: Caspase 7 (H69, H209, CCD8 treatment with etoposide)

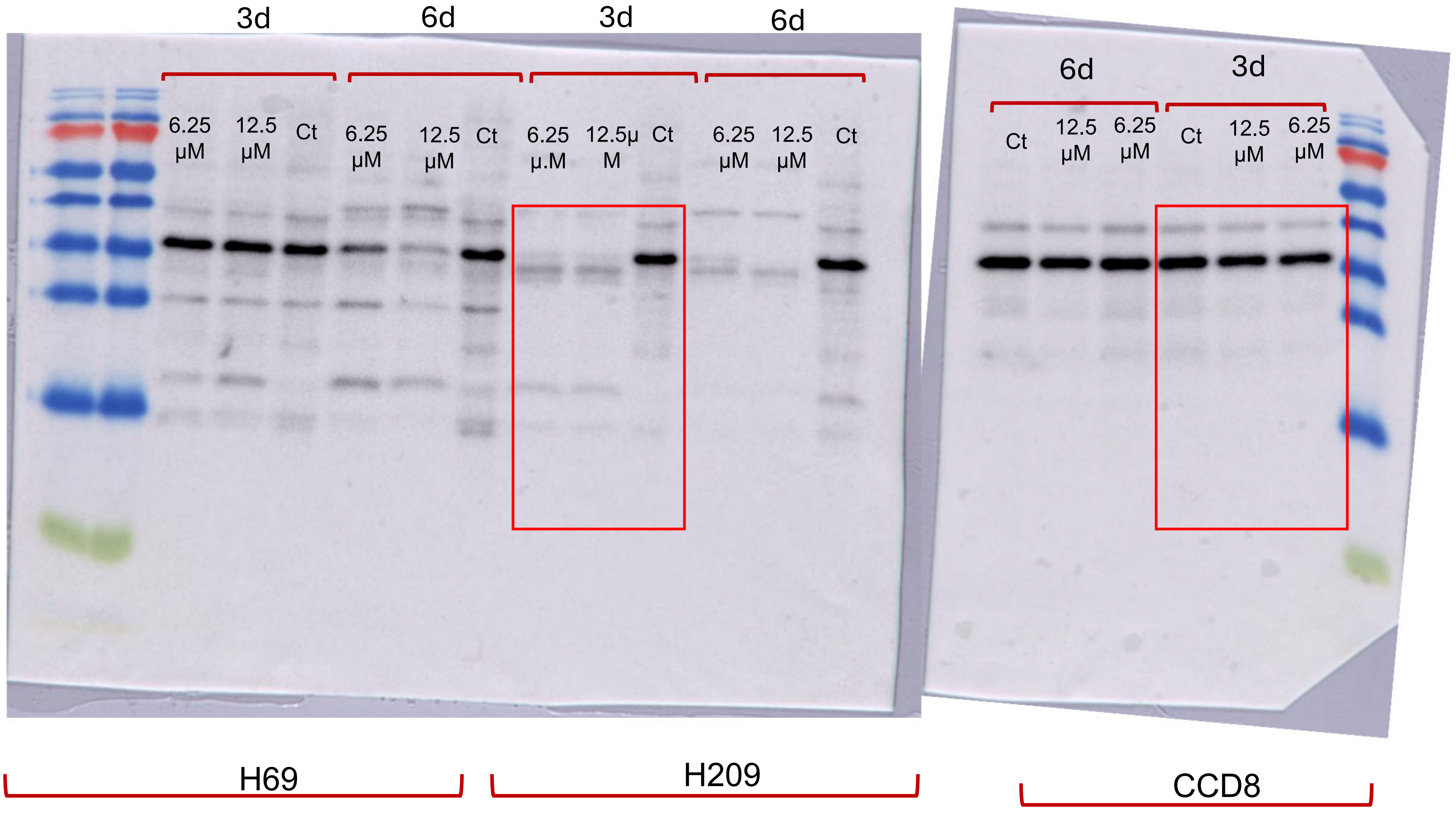

Supplemental Fig 1E: Caspase 3 (H69, H209, CCD8 treatment with cisplatin)

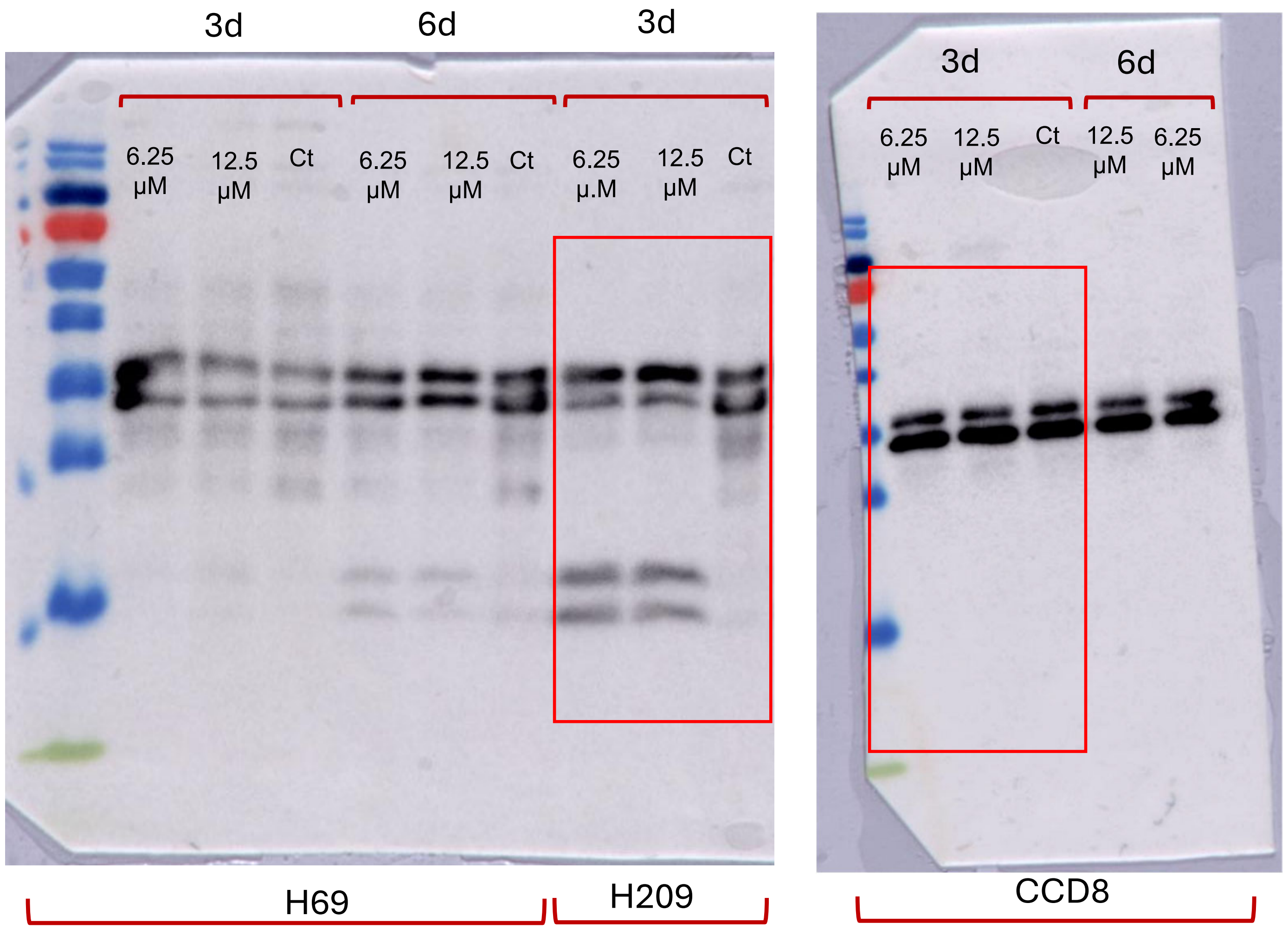

Supplemental Fig 1E: Caspase 7 (H69, H209, CCD8 treatment with cisplatin)

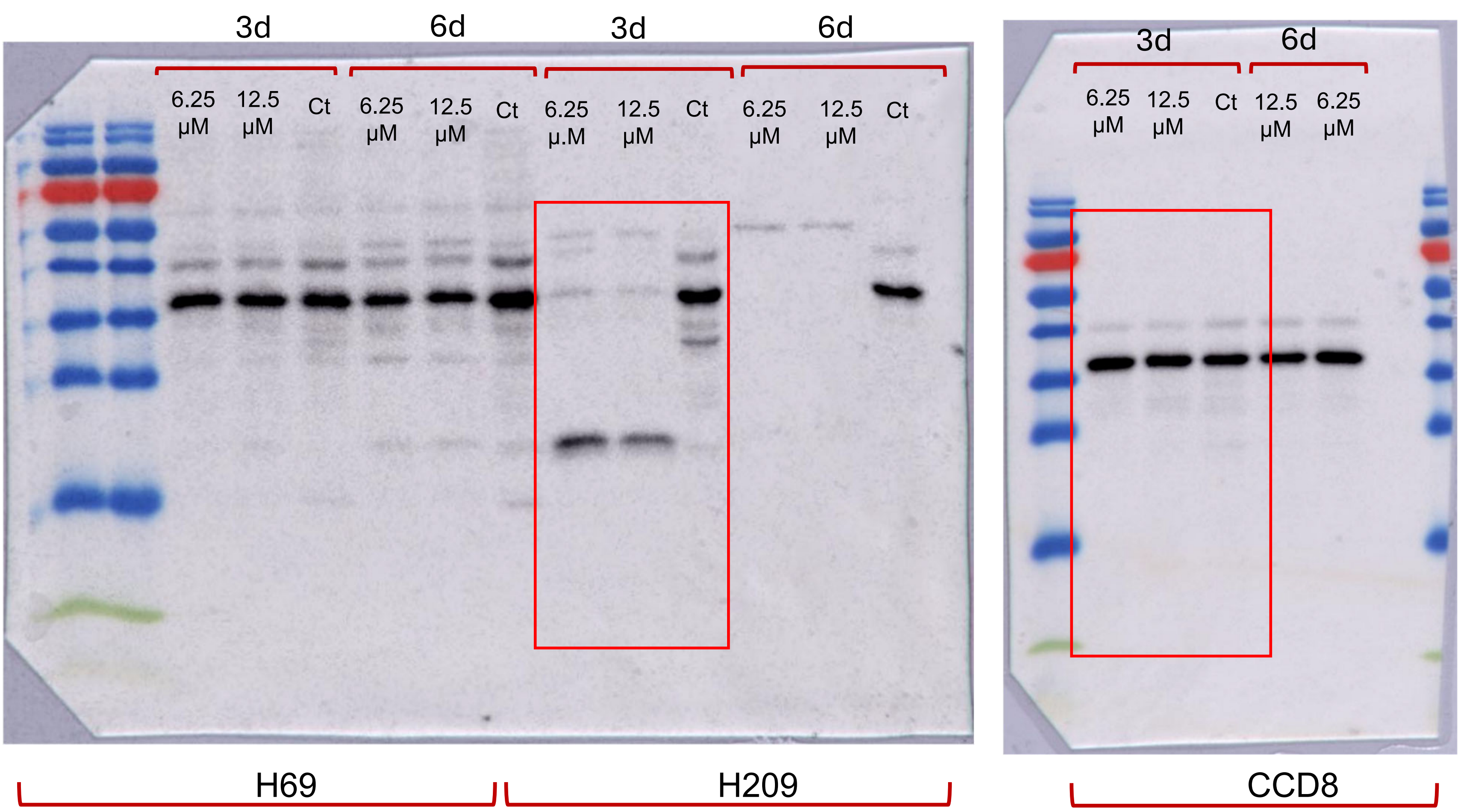

Figure 2C: Caspase 7 in H209 alone and H209 with CCD8 treated with Etoposide

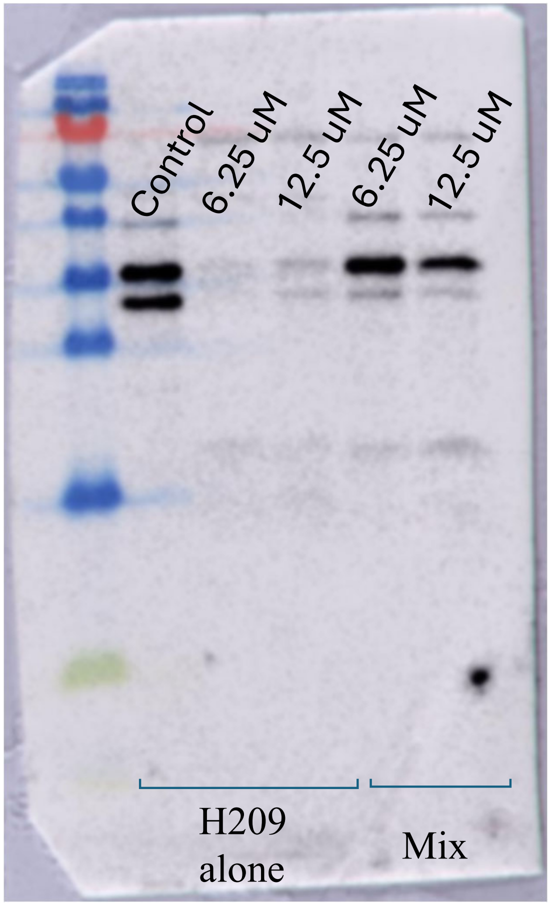

Fig 2D: Treatment with or without Verteporfin- caspase 3

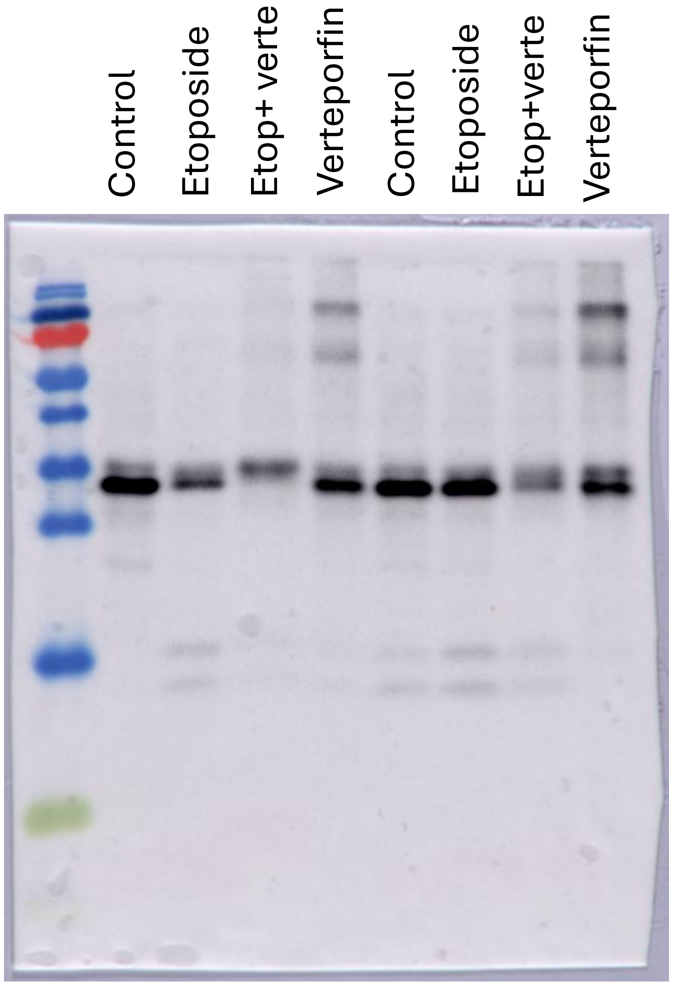

Fig 2E: Caspase 3- H209 grown in different conditioned media and treated with etoposide

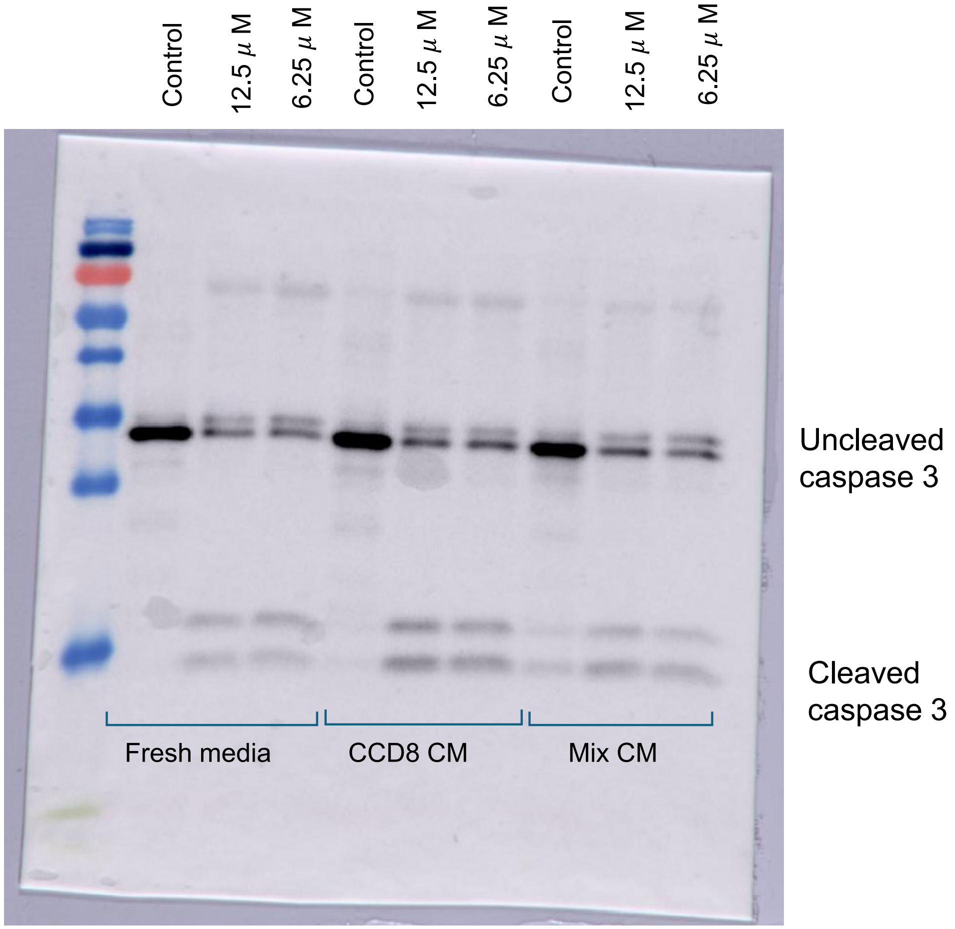

Supplemental Fig 2: Caspase 7- H209 grown in different conditioned media and treated with etoposide

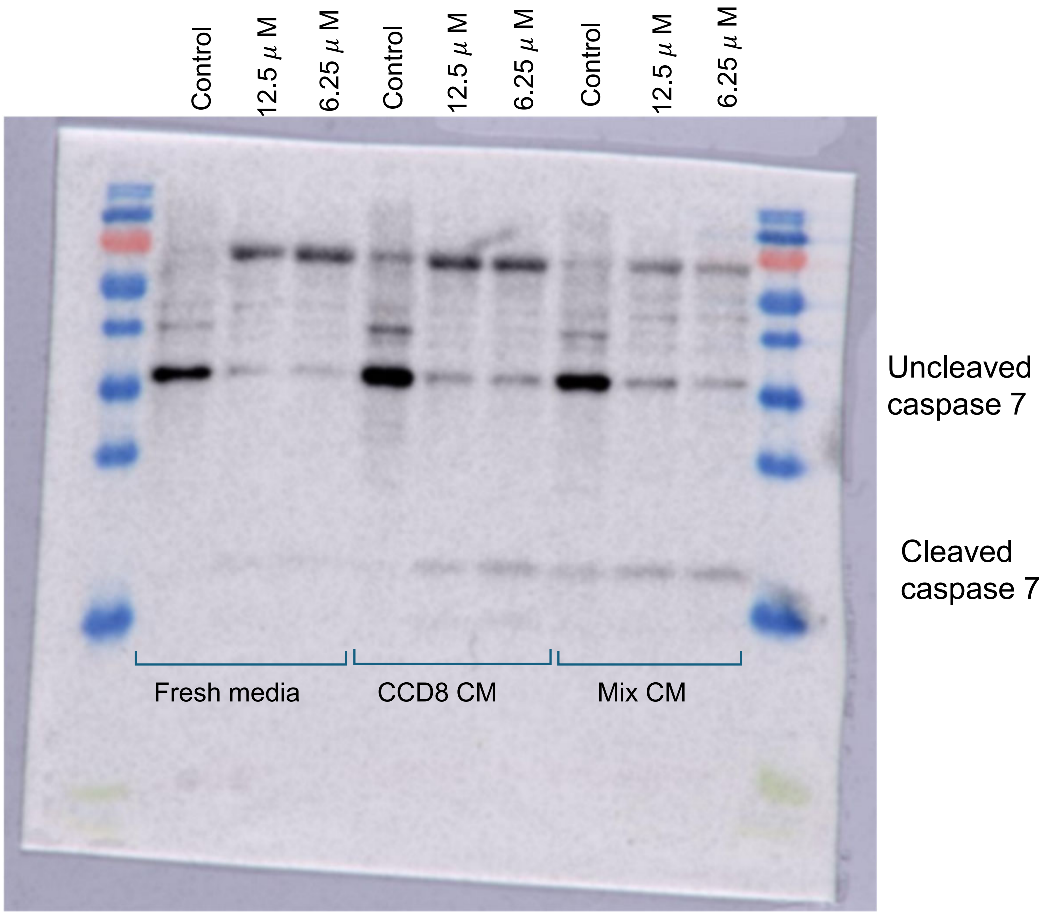

Fig 5A and 5B: Caspase 7- Drugs treatment of H209 and CCD8 for 36 hours

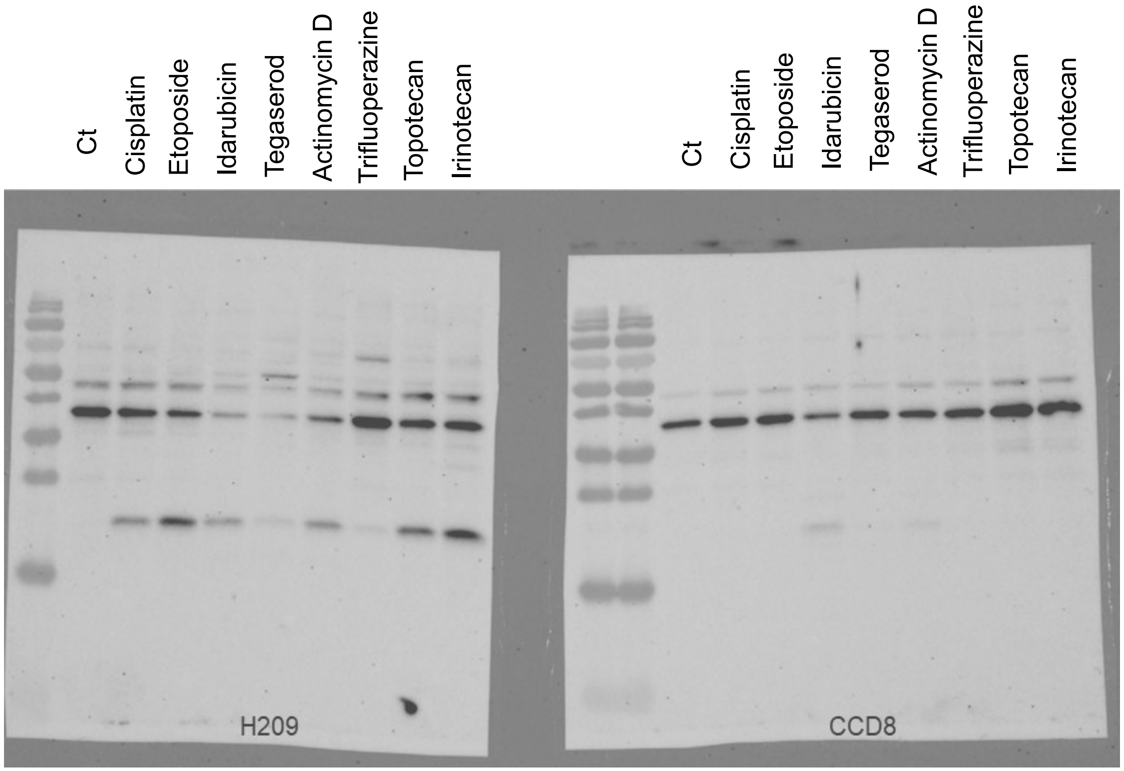

Fig 5C: Idarubicin dose response in H209 alone vs H209 co-cultured with CCD8

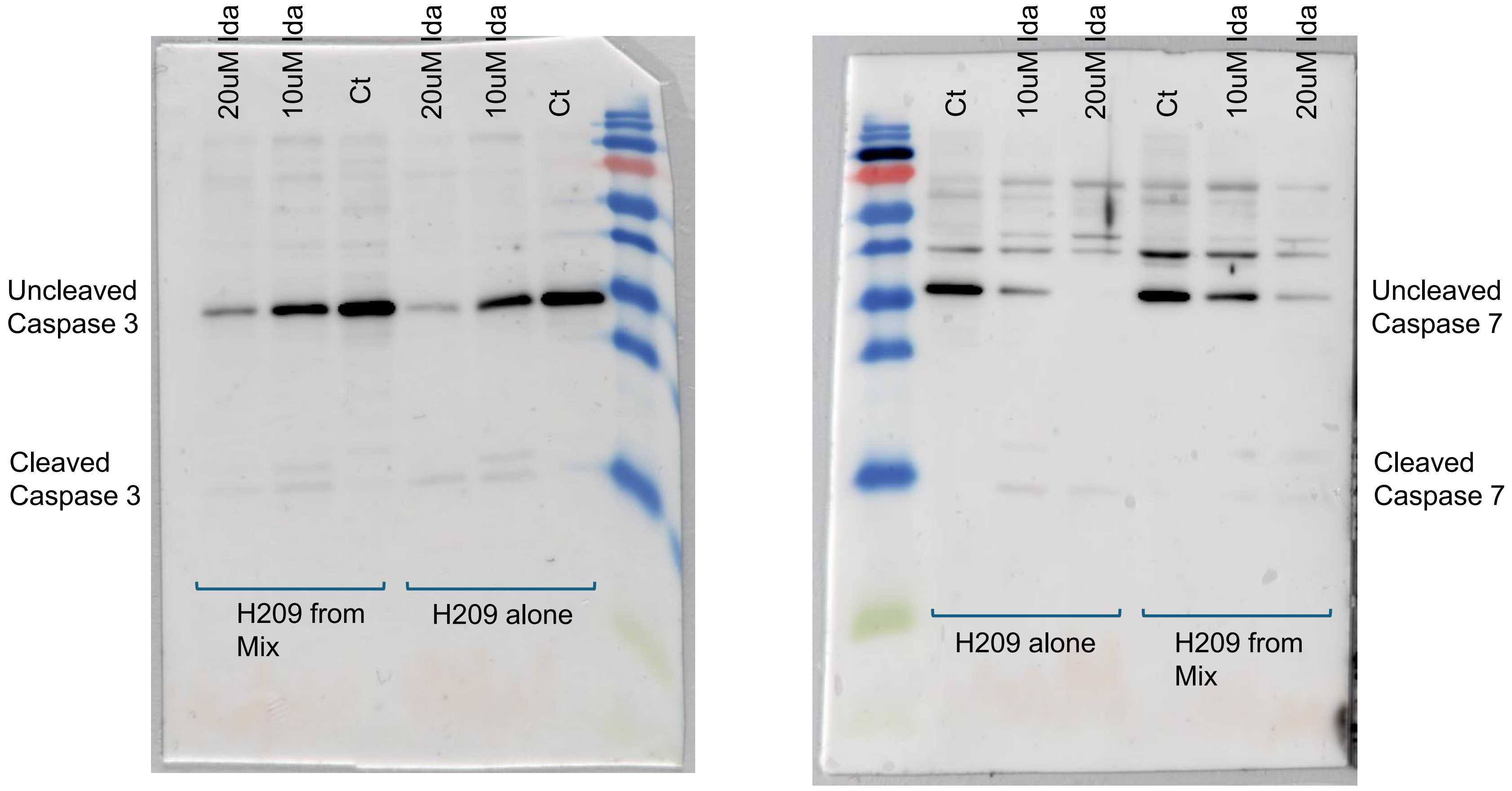

Fig 5D: H209 alone and from mix treated with Etoposide and Idarubicin for 48 hours

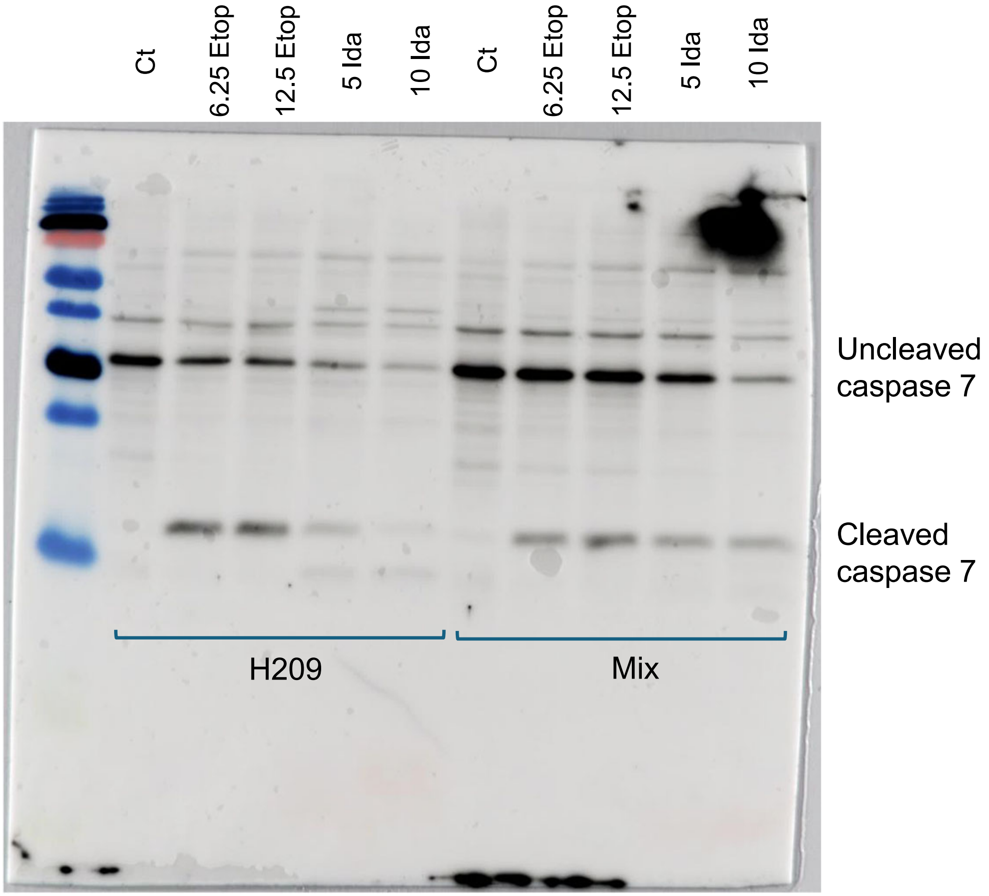
